# Transcriptome and metabolome analysis of crGART, a novel cell model of de novo purine synthesis deficiency: Alterations in CD36 expression and activity

**DOI:** 10.1101/2020.06.23.167924

**Authors:** Randall C Mazzarino, Veronika Baresova, Marie Zikánová, Nathan Duval, Terry G. Wilkinson, David Patterson, Guido N. Vacano

## Abstract

In humans, GART [phosphoribosylglycinamide formyltransferase (EC 2.1.2.2) / phosphoribosylglycinamide synthetase (EC 6.3.4.13) / phosphoribosylaminoimidazole synthetase (EC 6.3.3.1)] is a trifunctional protein which catalyzes the second, third, and fifth reactions of the ten step de novo purine synthesis (DNPS) pathway. The second step of DNPS is conversion of phosphoribosylamine (5-PRA) to glycineamide ribonucleotide (GAR). 5-PRA is extremely unstable under physiological conditions and is unlikely to accumulate in the absence of GART activity. Recently, a HeLa cell line null mutant for GART was constructed via CRISPR-Cas9 mutagenesis. This cell line, crGART, is an important cellular model of DNPS inactivation that does not accumulate DNPS pathway intermediates. In the current study, we characterized the crGART versus HeLa transcriptomes in purine-supplemented and purine-depleted growth conditions. We observed multiple transcriptome changes and discuss pathways and ontologies particularly relevant to Alzheimer disease and Down syndrome. We also report initial analysis of the Cluster of Differentiation (CD36) gene, which is highly expressed in crGART versus HeLa.

## 1. Introduction

*De novo* purine biosynthesis (DNPS) is one of the oldest and most fundamental biochemical pathways [1]. In mammals, starting with phosphoribosyl pyrophosphate (PRPP), the six enzyme, ten step pathway produces inosine monophosphate (IMP), which is subsequently converted to guanosine monophosphate (GMP) or adenosine monophosphate (AMP) via two more enzymatic reactions (Figure 1). Purines are critical as building blocks and carriers of genetic information in the form of RNA and DNA, intra and intercellular signaling molecules, energy currency, and substrates and co-enzymes. While salvage pathways can produce purine nucleoside monophosphates from free purine bases and PRPP, ultimately, all purines are produced by DNPS. DNPS is upregulated at the G1/S phase and is critical during cellular division [2], most likely to supply purines for DNA replication and elevated RNA transcription. Given its importance in cellular division, and that supply/transport of free purine bases across placental membranes is inefficient [3,4], DNPS is critical in mammalian development, including embryonic development. In mammals, the trifunctional GART enzyme catalyzes steps 2, 3, and 5 of DNPS

**Figure 1.**
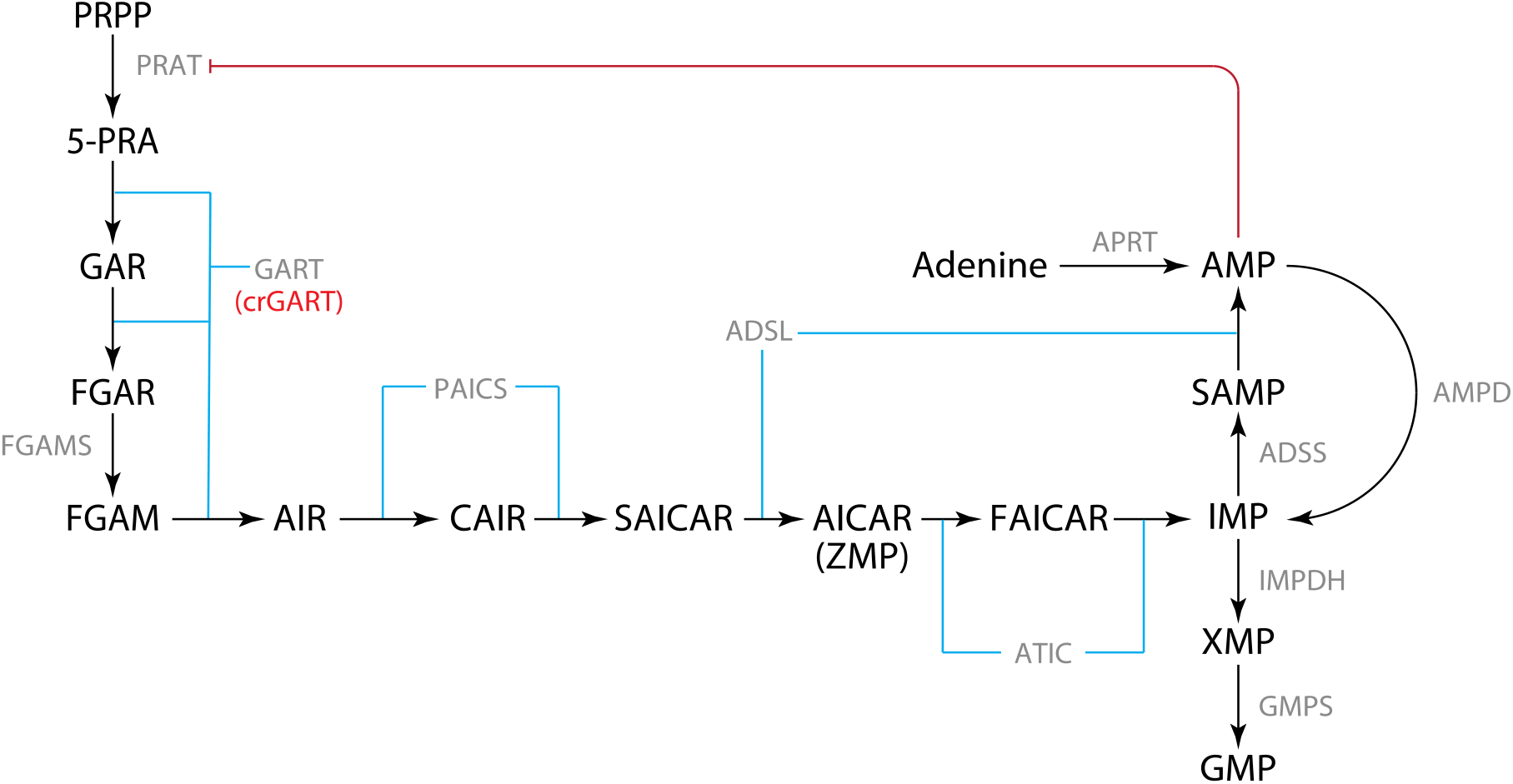
De novo purine synthesis (DNPS) pathway. DNPS mediates the conversion of PRPP to IMP. IMP is subsequently converted to AMP or GMP. The HeLa GART KO, crGART, is indicated.

(Figure 1). The human gene is located on Hsa21 and is therefore present in three copies in Down syndrome (DS, Trisomy 21), the most common genetic cause of intellectual disability in humans. Triplication of the GART gene has been hypothesized to be related to the pathology associated with DS [5]. GART expression is also altered in certain cancers [6,7] and is likely important for neurological metabolism and development [8,9].

The GART gene also encodes a monofunctional GARS protein via alternative transcription [5]. This alternative transcript includes a polyadenylation signal located in the intron separating the last GARS domain exon from the first AIRS domain exon. In human, mouse and Drosophila, this transcript has an in-frame TAA stop codon which is part of the 5′ donor splice site. This gene organization is not found in bacteria, lower eukaryotes, or plants. The function of GARS or the reason for this conservation of the gene structure confined to animals is not known. Characterization of the crGART transcriptome should allow identification of pathways in which GART plays a regulatory role and a better understanding of the GART gene and present an opportunity to assess the role of monofunctional GARS.

Functional mutations in DNPS genes are extremely rare in humans and have so far been limited to the adenylosuccinate lyase (ADSL), 5-aminoimidazole-4-carboxamide ribonucleotide formyltransferase / inosine monophosphate cyclohydrolase (ATIC), and phosphoribosylaminoimidazole carboxylase / phosphoribosylaminoimidazole succinocarboxamide synthetase (PAICS) genes. To date, fewer than 100 patients have been identified with ADSL deficiency. Until quite recently, only one patient had been identified with AICA-ribosiduria (ATIC deficiency), but now, three more individuals (two are siblings) have been identified [10]. Finally, two individuals (siblings) have been identified with PAICS deficiency [11]. ADSL deficiency is a spectrum disorder with three generalized classes: neonatal fatal, severe, or mild to moderate. Features of ADSL deficiency include seizures, psychomotor retardation, respiratory failure, and craniofacial abnormalities [12]. AICA-ribosiduria is characterized by mental retardation, blindness, epilepsy, and craniofacial and body dysmorphic features [6,9]. The siblings identified with PAICS deficiency died within three days of birth and exhibited craniofacial abnormalities and body dysmorphic features [11]. The majority of these mutations are amino acid point mutations ultimately resulting in decreased enzymatic activity. Thus far, no functional mutations in GART have been reported. Taken together, these data suggest that defects in DNPS are usually embryonic fatal.

DNPS nulls for ADSL, ATIC, and GART were recently generated in HeLa cells via CRISPR-Cas9 induced mutagenesis [13]. The ADSL and ATIC mutants (crADSL and crATIC) accumulate DNPS pathway intermediates when cultured in purine free media (the DNPS pathway is up-regulated in the absence of exogenous purines). These intermediates, SAICAR and ZMP respectively, are regulators of transcription, and we have presented evidence for transcriptional regulation via DNPS deficiency and intermediate accumulation ([14] and [15]). We were unable to detect intermediate accumulation in the GART mutant (crGART). This is expected since the initial substrate for GART is phosphoribosylamine (5-PRA), which is extremely unstable under physiological conditions and hydrolyses to ribose-5 phosphate; half-life 5 seconds [16]. We hypothesize that changes in transcription due to GART knockout in this cell line are due to the deficit in DNPS, and not intermediate accumulation. To evaluate the crGART transcriptome, we employed RNA-seq to compare the crGART and HeLa transcriptomes in adenine-supplemented and adenine-depleted conditions. Our results indicate that GART null cells have a large number of differentially expressed genes (DEG) and may have an important role in embryogenesis, neural development, and perhaps special relevance to Alzheimer’s disease (AD) and Down syndrome (DS). We focus on one DEG, Cluster of Differentiation 36 (CD36), which encodes a scavenger receptor that functions as a master regulator of long chain fatty acid (LCFA) metabolism. CD36 has been implicated in a number of health conditions, such as obesity, dyslipidemia, atherosclerosis, and AD. Given GART triplication and purine dysregulation in DS and the role of CD36 in conditions reminiscent of DS, we hypothesize that CD36 may have important roles in DS and associated conditions.

## 2. Materials and Methods

Methods were performed as previously described ([14] and [15]), except as noted.

### 2.1 Cell culture

Briefly, cells were maintained on 60 mm TPP (Techno Plastic Products, AG, Switzerland) plates in Dulbecco’s Modified Eagle Medium (DMEM, Gibco) supplemented with 10% fetal calf serum (FCS), 30 μM adenine, and normocin (Invivogen).

At two and one days prior to starvation induction, complete medium was replaced. Ten to twelve hours prior to adenine starvation, medium was changed to DMEM 10% FCS with 100 μM adenine. This concentration completely inhibits DNPS [17] and intermediate accumulation [14,15,18]. At ~50–70% confluence, medium was changed to DMEM supplemented with 10% fetal calf macroserum (FCM), normocin, with or without 100 μM adenine for 10 hour incubation. FCM is FCS dialyzed against saline using a 3.5 kDa molecular weight cutoff to deplete purines present in FCS.

### 2.2. HPLC analysis to detect metabolite accumulation

HPLC-EC analysis was performed as previously described ([14,18] and [15]). No DNPS intermediates were detected in adenine-supplemented or -starved cells.

### 2.3. RNA-seq

RNA-seq was performed as previously described ([14] and [15]).

### 2.4. Data processing

RNA-seq sequence (FASTQ format by the Genomics and Microarray Core Facility at the University of Colorado, Denver) was aligned to the Ensembl98 [19] Homo_sapiens.GRCh38.98 transcriptome using salmon version 1.0.0 [20], then differential gene expression analysis was performed using DESeq2 version 1.26.0 [21] with independent hypothesis weighting (IHW) [22] and approximate posterior estimation for GLM coefficients (apeglm) [23], R version 3.5.3 [24]. Differentially expressed genes are listed in Supplementary Table 1 - DEGs.

### 2.5. ClueGO analysis

The Cytoscape (version 3.7.2) app ClueGO (version 2.5.5) [25] was employed for ontology and pathway analysis of lists of differentially expressed genes (DEGs). A list of 300 DEGs with the top 150 most positive log2 and 150 most negative log2 values was used to query gene ontology (GO) databases (UniProt-GOA_08.01.2020 for Biological process, Cellular component, and Molecular function) and Reactome databases (Pathway_08.01.2020 and Reactions_08.01.2020). See Supplementary Figures 1-5 and Supplementary Table 2 - ClueGO.

### 2.6. qPCR validation of DEGs

The RNA-seq analysis was validated by qPCR of six DEGs covering a wide range of RNA-seq log2 fold differences (similar to previously described [12]). Samples were collected and processed approximately one year after collection and processing of samples for RNA-seq. Detection and amplification was performed with an Applied Biosystems QuantStudio7 Flex System and SsoAdvanced Sybr Green (BioRad cat #1725270). Candidate gene primers were obtained from IDT PrimeTime service: ALPP (Hs.PT.56a.38602874.g), DPYSL3 (Hs.PT.58.39796068), GATA3 (Hs.PT.58.19431110), IQGAP2 (Hs.PT.58.28018594), OASL (Hs.PT.58.50426392), TWIST1 (Hs.PT.58.18940950) and ACTB (Hs.PT.39a.22214847). Ct values and RNA-seq log2 values were normalized to ACTB (β-actin).

### 2.7. PANTHER analysis

PANTHER Overrepresentation Test analysis via the Gene Ontology Consortium portal (https://www.geneontology.com) was performed to confirm and augment the ClueGO results. Unlike ClueGO, PANTHER is capable of using large gene lists. Three gene lists were used to query PANTHER.

The complete list of significant DEGs, a list of DEGs unique to the crGART versus WT comparison (i.e., not present in previous analyses of crADSL and crATIC), and a list of DEGs common to the three mutants compared to WT were used to query Gene Ontology biological process, cellular component, and molecular function. See Supplementary Table 3 - PANTHER, Results 3.5 and 3.6 and Supplementary Table 4 - PANTHER unique, and Supplementary Table 5 - PANTHER common. The “Test Type” option was set for “Fisher’s Exact” and “Correction” was set to “Calculate False Discovery Rate”. The GO ontology database release date was 2020-02-21.

### 2.8. GART Transfection

crGART cells were transfected with pCMV-GART-K1 [26] using Lipofectamine 2000 as per manufacturer’s protocol (Invitrogen). Transfectants were cultured in high glucose DMEM, 10% fetal calf serum (FCS) supplemented with normocin, Geneticin (ThermoFisher Scientific), and 30 μM adenine (3 × 10^−5^ M). Stable transfectants were selected for Geneticin resistance and then checked for purine (adenine) growth requirement.

### 2.9. Protein Isolation

Confluent cell cultures (60 mm plates) were rinsed 2X with 1X PBS, then 500 μl cell lysis buffer (1% Triton X-100, 20 mM Tris pH 7.5, 50 mM NaCl, 5mM MgCl_2_) was added. The plates were scraped and lysate was collected and transferred to 1.5 ml Eppendorf tubes. Lysates were centrifuged for 30 minutes at 16,000 x g. The supernatant was collected and protein concentration was measured by Bradford assay (Sigma).

### 2.10. Western Blot

Total HeLa, crATIC, crGART, and crGART/K1-GART (transfected cells) protein (30 μg/lane) was separated by SDS-PAGE (8-16% gradient tris-glycine gel) for 40 minutes at ~225 V. Gels were wet-blotted to nitrocellulose (buffer: 25 mM Tris, 200 mM glycine, 20% methanol) at 200 mA for 1 h. Blots were blocked 30 minutes in block solution (1% BSA, 0.1% Tween-20, 1X TBS, 0.01% sodium azide, pH 7.4) then incubated for 1.5 h at RT with either CD36 (Novus NB400-144, 1:500 dilution) or PPARγ (Cell Signaling 2430S, 1:250 dilution) and GAPDH (Abcam ab181602, 1:10,000 dilution) rabbit primary antibodies in primary Ab solution (5% BSA, 0.1% Tween-20, 0.01% sodium azide, 1X TBS, pH 7.4). Blots were washed 3X for 5 minutes in 1X TBS. Blots were then incubated in goat anti-rabbit IgG-AP conjugate (Bio-Rad, 170-6518) diluted 1:10,000 in block solution for 1 hour. Three more washes (5 minutes each in 1X TBS) were performed, then blots were treated with CDP-Star® Substrate with Nitro-Block-II™ Enhancer (Applied Biosystems, T2218) for five minutes and visualized using a Bio-Rad Chemidoc Imager.

### 2.11. Metabolon metabolomic analysis

Cells were grown and starvation induced in 100 mm TPP cell culture dishes as described above. Cells were harvested according to Metabolon’s prescribed protocol. Briefly, cells were washed in 1X PBS and treated with Detachin (Genlantis) to release cells from the plate. Cells were transferred to 15 ml conical tubes and 10 ml of experimental condition media was added. Cells were pelleted at 1000 x g for 5 minutes, the pellet was aspirated, resuspended in 1 X PBS and pelleted again, aspirated, and stored at −80°C until shipment to Metabolon on dry ice. Samples were analyzed by Metabolon for metabolomic analysis using ultra high-performance liquid chromatography/tandem accurate mass spectrometry (UHPLC/MS/MS). Peak detection and identification, determination of relative concentrations, and statistical analysis were provided by Metabolon (http://www.metabolon.com).

## 3. Results

### 3.1. Adenine is required for proliferative growth of crGART

Given that GART catalyzes three of the ten steps in the DNPS pathway, we hypothesized that crGART cells would exhibit proliferative arrest in adenine-deprived conditions. HeLa and crGART cells were cultured in complete-serum media overnight and then subsequently cultured in media supplemented with dialyzed-serum (FCM) with or without supplemental adenine. HeLa cells showed proliferative growth in both media conditions while crGART showed proliferative growth in only adenine-supplemented conditions (Figure 2).

**Figure 2.**
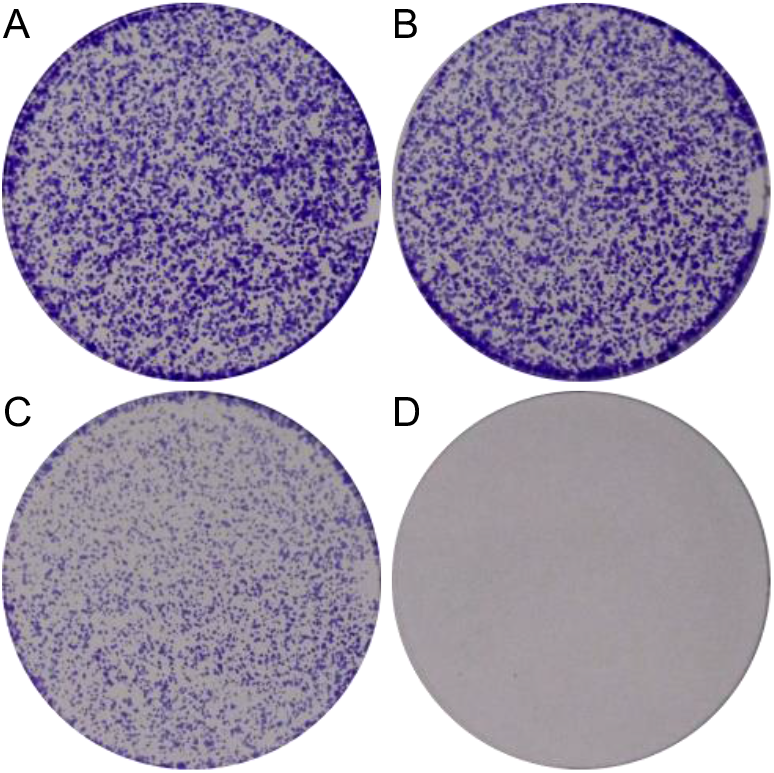
crGART requires adenine for growth. HeLa (A,B) and crGART (C,D) cells were plated and cultured in 30 μM adenine-supplemented (A,C) or adenine-depleted (B,D) media. Plates were stained with crystal violet.

### 3.2. DNPS intermediates were not detected in adenine starved crGART cells

Next, we assessed whether the crGART cells accumulate any detectable DNPS intermediates. We demonstrated previously that crADSL [14] and crATIC [15] cells accumulate substrate when cultured in adenine free media. crGART cells were cultured as previously described and metabolites extracted. HPLC-EC analysis did not indicate accumulation of any DNPS intermediates (data not shown). This is consistent with previous work demonstrating that 5-PRA is highly unstable and breaks down in approximately five seconds under cellular conditions to ribose-5 phosphate [12,14] [27]. This suggests that crGART is likely useful as a model of general DNPS deficiency.

### 3.3. Transcriptome analysis of crGART versus HeLa identified differentially expressed genes

To investigate the effect of GART KO on the transcriptome, we compared the crGART and HeLa transcriptomes after culture in adenine-supplemented and adenine-depleted conditions. 4218 genes were significantly differentially expressed by cell type. This represents a log(2)counts crGART: HeLa fold change range of −13.94 to 10.93 (Figure 3A). For ClueGO analysis, we used the “top 300” DEG list (see Materials and Methods 2.5), which represents a log(2)counts fold change range of −13.74 to −7.67 and 5.83 to 10.93 (Figure 3B). Positive values indicate enrichment in crGART and negative log2 values indicate enrichment in HeLa. Principal component analysis shows clustering by cell type and supplementation (Figure 4).

**Figure 3.**
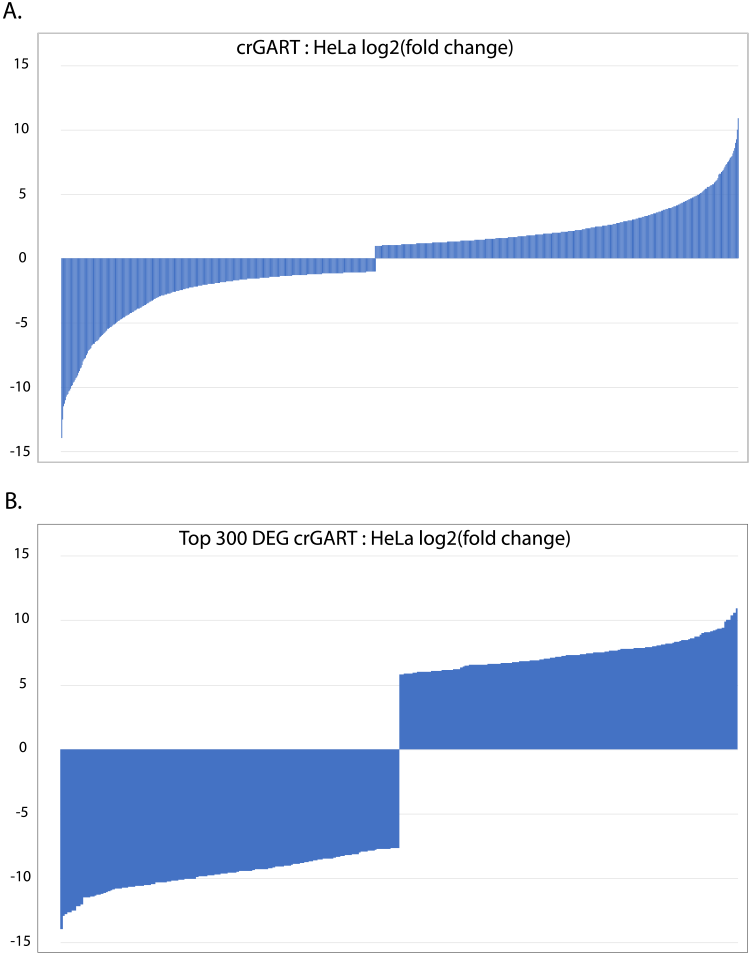
Plot of log(2) values crGART: HeLa DEG fold change. A. All significant DEGs B. Significant DEGs in the “top 300” list (see Methods).

**Figure 4.**
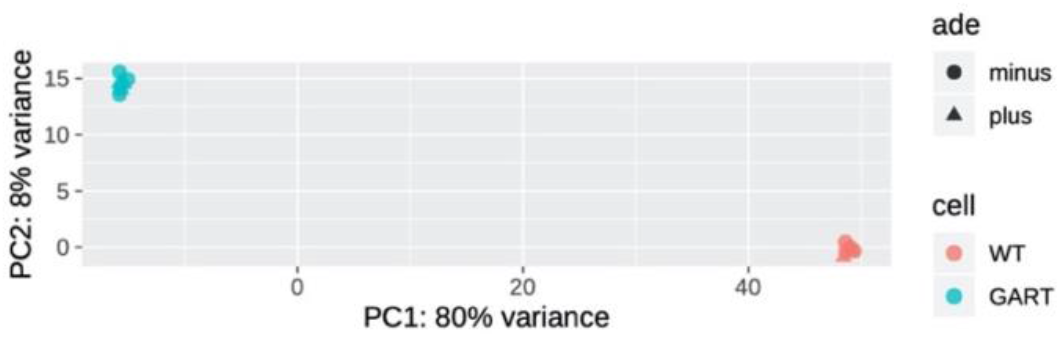
Principal component analysis of crGART and HeLa replicates. PCA shows a robust difference by cell type and a much smaller difference by adenine supplementation.

### 3.4. CD36

One of the largest magnitude DEGs we obtained was for the CD36 gene. Transcript levels for the Cluster of Differentiation 36 (CD36) gene are highly elevated in crGART versus HeLa (Figure 5A, Supplementary Table 1 - DEGs) and in crADSL [14] and crATIC [15], which suggests that DNPS regulates expression of the gene. Additionally, transcript levels for the Peroxisome Proliferator Activated Receptor γ (PPARG) gene are elevated in the mutant cell lines (Figure 5C).

**Figure 5.**
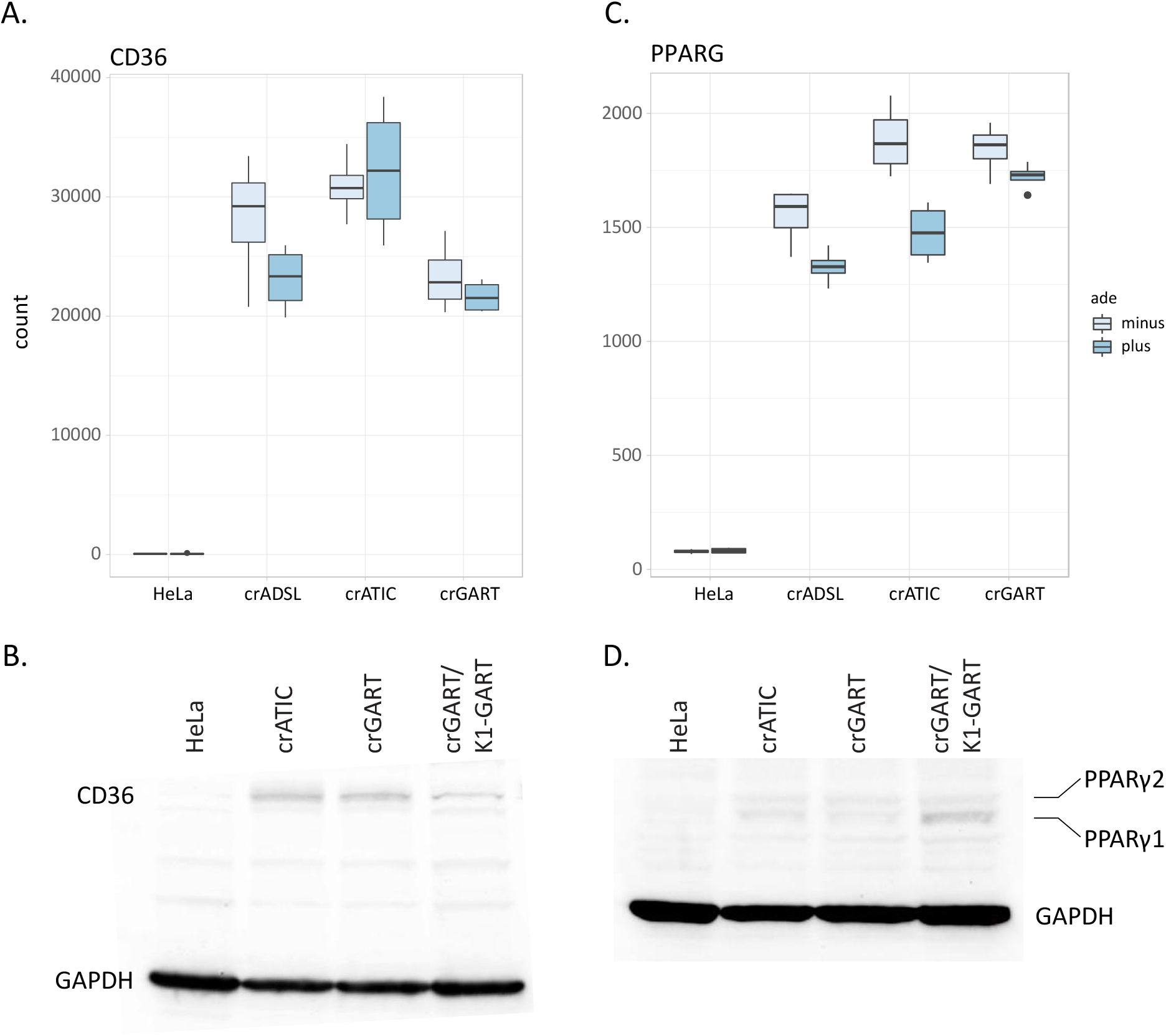
**A**. Box plot showing CD36 RNA-seq transcript levels in HeLa and DNPS mutants **B**. CD36 immunoblot **C**. Box plot showing PPARG RNA-seq transcript levels **D**. PPARγ immunoblot. The predicted MW of CD36 is 75 kDa, and the predicted MW of PPARγ1 and PPARγ2 are 53 and 57 kDa. The MW of GAPDH is 36 kDa.

To evaluate CD36 and PPARγ protein expression, we performed immunoblot analysis, which confirmed elevated expression in crATIC and crGART, and relative absence of the two proteins in HeLa (Figure 5B and 5D). Our results also show that elevated CD36 expression is partially rescued in crGART cells stably transfected with a CHO-K1 GART expression construct.

In a separate set of experiments, we conducted a metabolomic analysis of crGART and crGART stably transfected with a GART expression construct via the Metabolon Global Metabolic Panel analysis. In this analysis, 22 long-chain fatty acids were identified, and of these, about 15 are markedly elevated in the transfected cells (Figure 6, Supplementary Table 6 - Metabolon LCFA). This is especially interesting given that CD36 is an important regulator of long-chain fatty acid metabolism [28] and that CD36 expression is reduced in the transfected cells.

**Figure 6.**
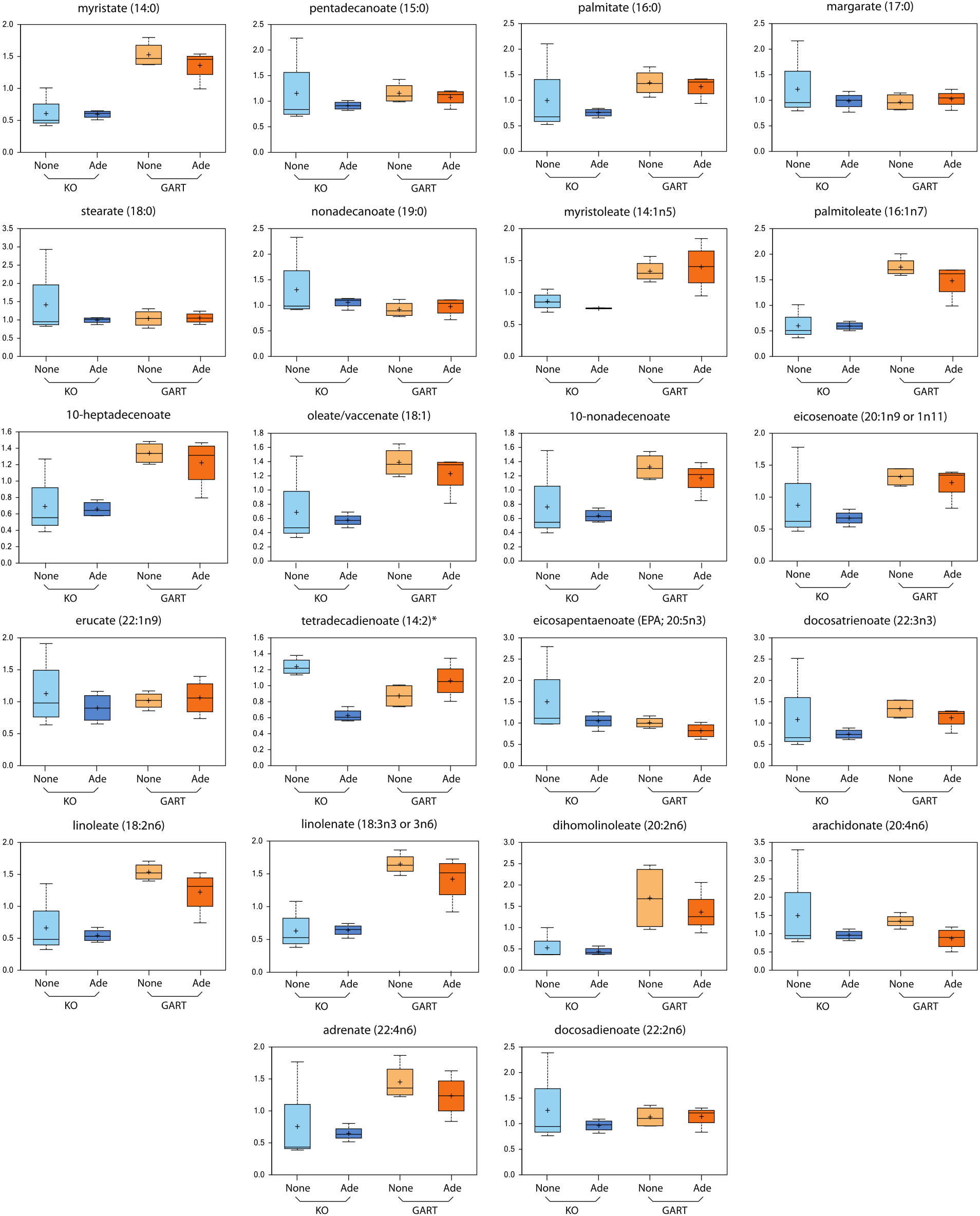
Box plots supplied by Metabolon, Inc. showing LCFA levels in crGART (KO) cells and transfected crGART (GART) cells. Ade indicates that cells were grown in purine supplemented media, None indicates purine starvation.

### 3.5. ClueGO analysis: Functional enrichment analysis employing Gene Ontology and Reactome databases

In order to assess what processes or pathways are affected by ablation of the GART gene, we performed GO and Reactome functional analysis of DEGs. We queried the Gene Ontology database (Biological process, Cellular component, and Molecular function) as well as the Reactome knowledgebase (Pathways and Reactions) for this analysis. We obtained a very large number of terms but will focus only on terms of special interest to our laboratory (Figure 7). All data is available in supplementary material.

**Figure 7.**
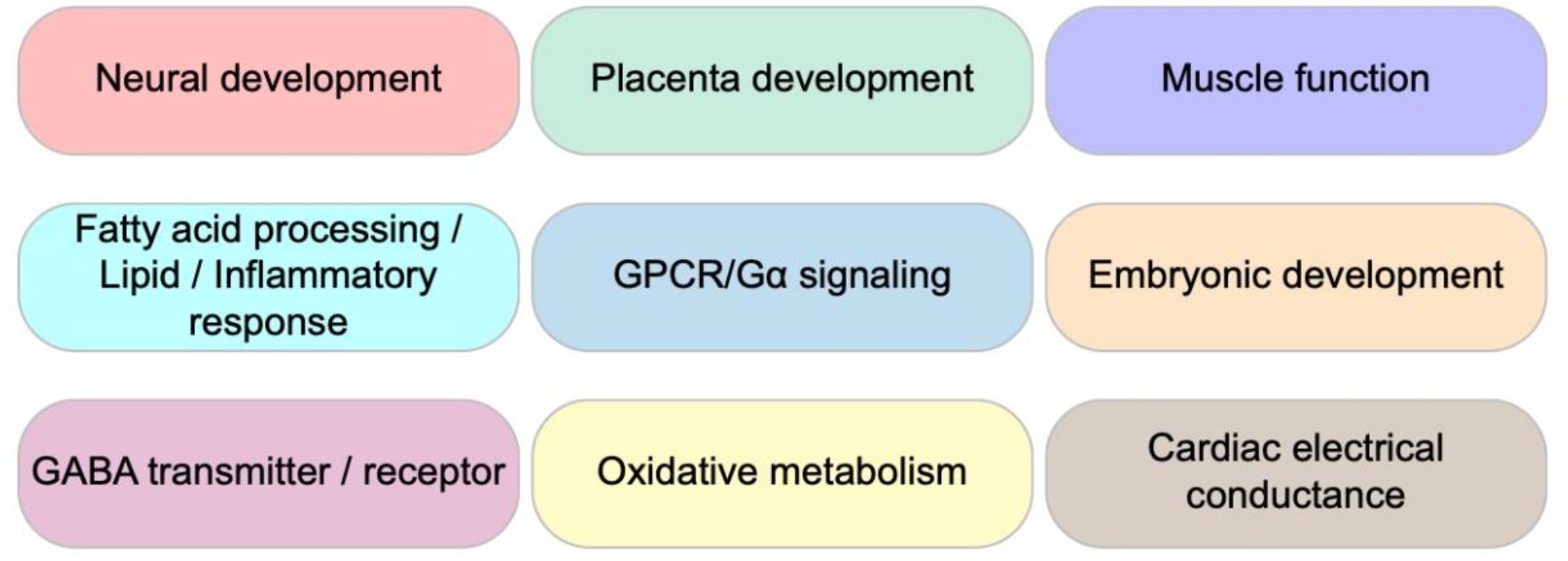
Prominent GO ontologies from ClueGO analysis

For the GO terms associated with Biological process, 70 GO groups were identified with 548 GO terms. Related GO groups were organized into neural function, development, muscle function, fatty acids, cardiac function, G-protein coupled activity, oxides, and lipids categories. Within neural function, associated terms include neuron migration, dendrites, axon, hippocampus, neuroblast proliferation, synapse, GABA, and catecholamines. DNPS is known to be important for neural function and diseases associated with DNPS exhibit strong neurological phenotypes. Interestingly, another category identified was development. Development includes disparate terms, ranging from organ morphogenesis related terms such as renal, pancreas, and prostate development, to mesenchyme morphogenesis, stem cell differentiation, alkaline phosphatase activity, blood vessel morphogenesis, epithelial cell differentiation, bone development, endoderm, etc. Placental development included labyrinthine development and female pregnancy. For muscle function terms, we obtained smooth muscle function, myotubes and sarcomere organization, vasoconstriction, actin filament movement, and striated and smooth muscle contraction. Cardiac function was also enriched, including sinoatrial and atrioventricular node function terms, as well as atria and ventricle development. Oxidation related terms such as nitric oxide and superoxide metabolism were found. Lipid terms included fatty acids, phospholipase activity, phosphatidyl metabolic process, PI3K, and inflammatory response centered around IL-1 and TNFα. Given the importance of purines in development and neurobiology, the observed enrichment in related terms was expected. However, enrichment in cardiac terms involving electrical conduction was unexpected and may suggest a novel line of inquiry. The enrichment for placental development terms was also unexpected and suggests a purine requirement early in development and that DNPS may play an important role in blastocyst formation.

For the GO terms associated with Cellular component, 12 groups were identified with 26 terms. Terms include voltage gated potassium channels, I-band and intercalated disc, gap junction, GABA synapse and various other synaptic membrane terms.

For the GO terms associated with Molecular function, 25 groups were identified with 54 terms. Term categories included neuronal and ion transmembrane function, development, phospholipid activity, cytokine, etc. In the neuronal category, neuropeptide, acetylcholine receptor, potassium ion gate channels, and neurotransmitter transmembrane transporter terms were prominent. For phospholipids, notable terms included lipase activity, phospholipase (A2, 1, and C) activity, phosphatidylcholine acylhydrolase activity, phosphatidylinositol 3 kinase, and nitric oxide synthase. Enrichment for Syndecan protein binding and G-protein coupled peptide receptor terms is consistent with the G-protein coupled activity category in the Biological process results above.

Reactome Pathways knowledgebase showed 14 groupings of 29 terms. Categories include diseases associated with O-glycosylation of proteins, peptide hormone metabolism, rhodopsin-like receptors, and GPCR and G-alpha signaling. Neuronal and development terms were identified, as well as TFAP-2 which is heavily involved in neural-crest development. Consistent with the GABA terms we obtained in Biological process, we identified GABA receptor and GABA B receptor terms. This suggests that neural deficits associated with DNPS or GART deficiency may be associated with alteration of GABAergic signaling.

Reactome Reactions knowledgebase showed 10 groupings of 24 terms. Term categories included proteoglycans and GPI anchors, GPCR, GEF, and Gi activated ligands. Keratin filaments was also noted.

ClueGO networks, pie charts, and ontology histograms are shown in Supplementary Figures 1-5.

### 3.6. Corroboration of Gene Ontology analysis via PANTHER analysis

PANTHER analysis with the complete list of significant DEGs was consistent with the ClueGO results and revealed new ontology terms such as DNA and RNA binding, polymerase, and transcription regulation (Supplementary Table 3 - PANTHER).

### 3.7. Comparison of crATIC, crADSL and crGART DEGs

To get a better sense of which DEGs are due specifically to ablation of the GART gene, we identified the significant DEGs unique to the crGART vs HeLa comparison. There were 1282 genes that changed significantly in the crGART vs HeLa comparison but did not change in the crADSL vs HeLa [14] and crATIC vs HeLa comparisons [15]. The list of genes was used to query PANTHER via the Gene Ontology web portal (www.geneontology.org). We obtained Biological process ontologies related to RNA splicing and mitochondrion organization (Supplementary Table 4 - PANTHER unique).

We also identified DEGs common to all three mutants compared to HeLa. PANTHER functional enrichment analysis of the list of 7745 genes returned numerous Biological process ontologies, including regulation of protein localization to plasma membrane, regulation of osteoblast differentiation, positive regulation of I-kappaB kinase/NF-kappaB signaling, heart morphogenesis, establishment of vesicle localization, Ras protein signal transduction, embryonic organ morphogenesis, and many others (Supplementary Table 5 - PANTHER common). This result illustrates the importance of DNPS in multiple aspects of cell development and function.

### 3.7. Validation of RNA-seq by qPCR of candidate genes

To confirm our RNA-seq results, we performed qPCR to measure transcript levels for ALPP, DPYSL3, GATA3, IQGAP2, OASL and TWIST1, and normalized to ACTB. These genes span a wide range of transcript levels. The transcript level results (Ct values) are consistent with the log(2)fold values for these genes from our RNA-seq analysis (Figure 8).

**Figure 8.**
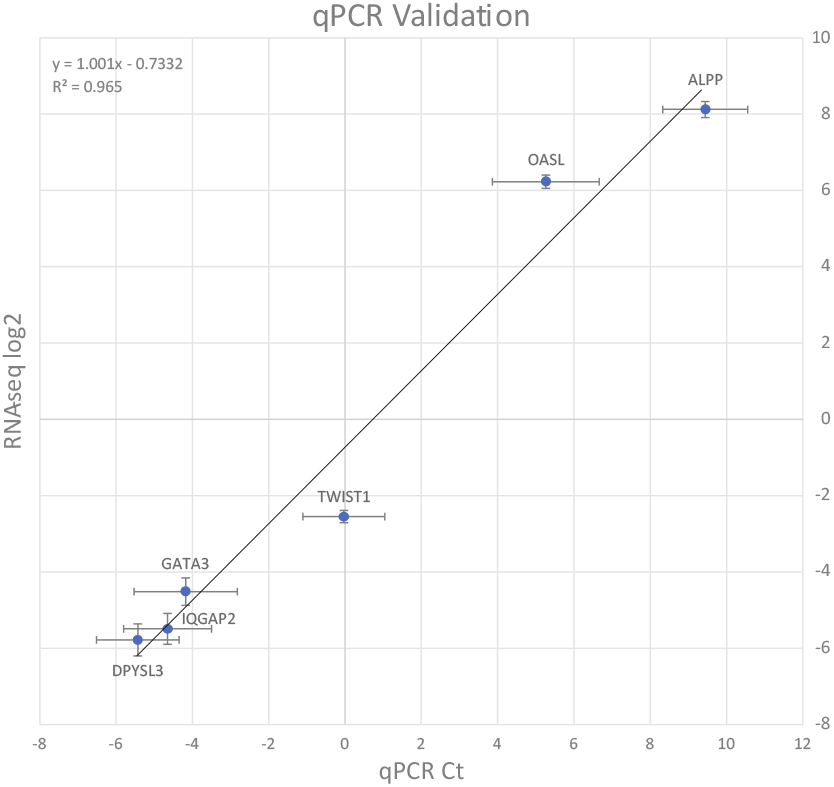
Validation of RNA-seq by qPCR. RNA-seq log2 values are plotted on the Y axis and qPCR Ct values are plotted on the X axis. Error bars indicate standard error.

## 4. Discussion

In this study, we evaluated the purine dietary requirement for crGART and performed HPLC-EC to detect accumulation of metabolites during purine starvation. We compared the crGART and HeLa transcriptomes via RNA-seq analysis in adenine-supplemented and adenine-depleted conditions.

The purpose of adenine supplementation is twofold: 1) to provide purines necessary for cell survival and 2) to eliminate DNPS [29,30], preventing intermediate accumulation. Adenine phosphoribosyltransferase (APRT) catalyzes the conversion of adenine and phosphoribosyl pyrophosphate (PRPP) to AMP and pyrophosphate (PPi). APRT is present in all mammalian tissues and is uniquely responsible for metabolic adenine salvage from dietary sources [31]. Once AMP has been produced, it can be converted to IMP by AMP deaminase (AMPD), and then converted to GMP by IMP dehydrogenase (IMPDH, converts IMP into XMP) and GMP synthase (converts XMP into GMP, see Figure 1) [32].

The GART gene encodes a trifunctional protein and also a monofunctional GARS protein via alternative transcription. The conversion of 5-PRA to GAR is the first reaction and is catalyzed by the GARS domain of the trifunctional protein [16]. As discussed previously, 5-PRA is extremely unstable under physiological conditions, and is unlikely to accumulate. Our results show that crGART requires purine (adenine) supplementation for proliferative growth, but apparently does not accumulate pathway intermediates during purine starvation (data not shown). In the crADSL and crATIC DNPS-KO models, metabolic substrates (SAICAR and ZMP respectively) readily accumulate during purine starvation and there is strong evidence that both substrates alter cellular processes [33–35]. Since the crGART line does not accumulate detectable DNPS metabolic intermediates during purine starvation, it is likely a useful model of strict DNPS deficiency.

The biological significance of the monofunctional GARS protein has not yet been elucidated. One hypothesis is that GARS may interact with PRAT to facilitate transfer of 5-PRA, the unstable product of PRAT, to GARS/GART thus increasing the rate of DNPS [12,32]. This would imply that lack of GARS would inhibit DNPS. The pCMV-GART-K1 expression construct that we used to transfect crGART encodes only the trifunctional GART gene, not the monofunctional GARS gene. The transfected cell line is able to synthesize levels of purines adequate for growth, but purine levels may be less adequate, or inadequate, for other DNPS functions. This would be consistent with the observation that the transfected crGART cell line does not exhibit fully restored inhibition of CD36 expression. Since the GART gene is trisomic in DS, GARS overexpression may play a role in elevated purine levels in DS. Other hypotheses exist, for example, the monofunctional GARS protein may have a role in release of metabolites from the *de novo* purine pathway so they can be used in other metabolic pathways. The crGART cell line should be invaluable for investigating the role of the monofunctional GARS protein.

We noted that the CD36 gene is highly expressed in all three DNPS-KO transcriptomes that we have characterized (crADSL, crATIC, and crGART), with average log2(fold change) value of about 8.5 (Figure 5A). We have shown that CD36 protein expression is elevated in crATIC and crGART versus HeLa (Figure 5B) and that protein expression is partially rescued (reduced) in crGART cells stably transfected with a CHO-K1 GART expression construct. Finally, we show that numerous LCFA are elevated in transfected crGART, which may be connected to reduced CD36 expression in these cells. CD36 is involved in fatty acid uptake and metabolism [28], inflammation cascade [36], angiogenesis, apoptosis, thrombosis, atherosclerosis, Alzheimer’s disease, and insulin resistance [37]. Interestingly, many of these phenotypes are seen in DS and associated conditions. We hypothesize that CD36 is regulated by DNPS and is dysregulated in DS. This hypothesis is supported by an RNA-seq analysis of age and sex matched fibroblast cell lines derived from people with DS and normosomic controls showing that CD36 expression is reduced two-fold in the DS lines (see supplemental table from [38]). As such, reduced CD36 expression/elevated LCFA levels may play an important role in DS phenotypes and associated conditions (Figure 9).

**Figure 9.**
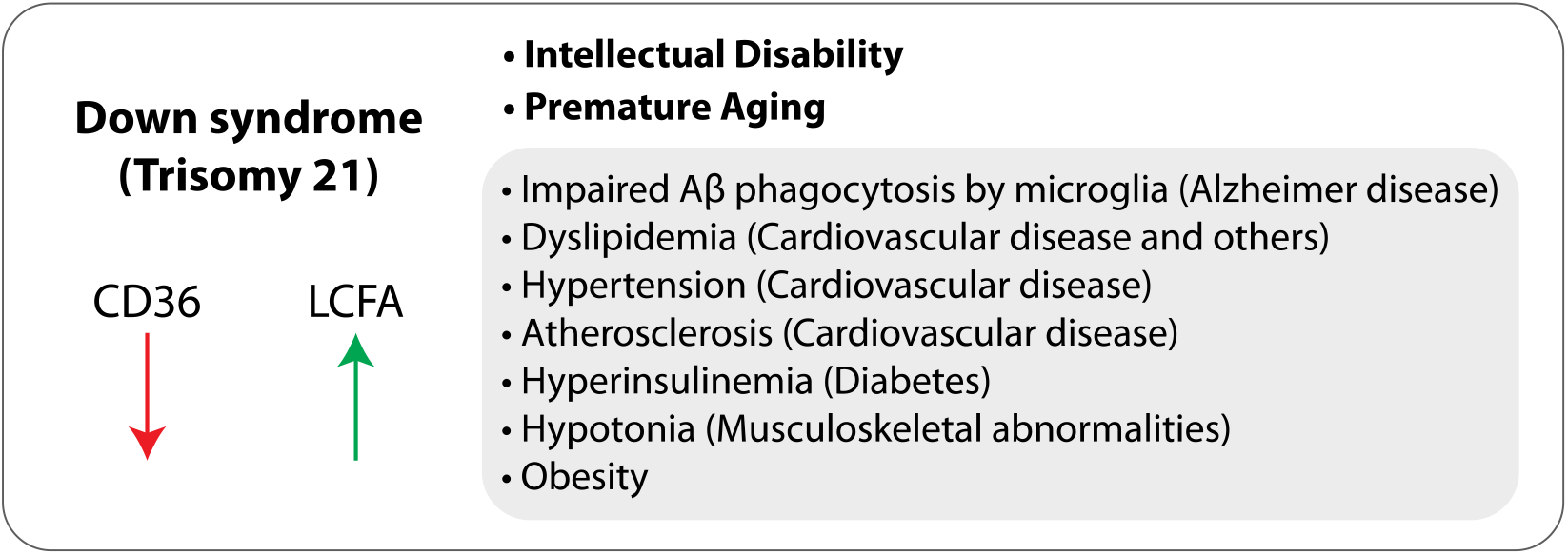
Suggested role of CD36 in Down syndrome. We hypothesize that reduced CD36 expression and increased LCFA levels may be important in intellectual disability and premature aging, as well as co-morbidities characteristic of DS.

The PPARG gene is also highly expressed in all three DNPS-KO transcriptomes (Figure 5C) and protein expression is elevated in crATIC and crGART (Figure 5D). PPARG encodes PPARγ, an inducible transcription factor that is considered a master regulator in lipid metabolism, with roles in fat, carbohydrate, and general energy metabolism, as well as insulin sensitivity, cell proliferation and differentiation, inflammation, and cancer [39]. A CD36-PPARγ axis regulates fatty acid storage, triglyceride synthesis, glucose uptake, and fatty acid metabolism [40].

In addition to CD36 and PPARG, our RNA-seq analysis led to identification of numerous DEGs and gene ontologies from ClueGO functional enrichment analysis. Many of these were consistent with our previous analyses of the crADSL and crATIC mutants and are likely to be due to DNPS deficiency. These results support the hypothesis that DNPS is essential in development and that alterations in DNPS and intermediate metabolite accumulation both regulate the transcriptome. This is consistent with previous work demonstrating that DNPS is upregulated in the G1/S phase cell cycle interface [3,4], is critical in embryogenesis which is marked by rapid cellular division, and that purines are essential in development [8] especially in neural development [41].

We also obtained DEGs and gene ontologies potentially relevant to DS. This is perhaps unsurprising given that 1) the GART gene is located on Hsa21 and is triplicated in DS, 2) GART expression is dysregulated in DS and 3) purine levels are also dysregulated in DS [42]. In addition, although the current work involves a GART null model, it is likely that at least some pathways affected by absence of GART are also affected by increase in GART, although the relationship is unlikely to be symmetrical or straightforward. Ontologies include terms relevant to placental development, neural development and cognition, cardiac development, and Alzheimer disease.

People with DS are at elevated risk for developmental as well as aging related disorders. In addition to intellectual disability, common disorders include hypotonia [43], congenital heart malformation, disease of pulmonary circulation, cardiac arrest, hypotension, infantile spasm, epilepsy, OSA (sleeping disorders), dementia, hypothyroidism, and obesity [44]. Our transcriptomic analysis revealed multiple terms potentially relating GART and DNPS to these disorders. Ontological groupings show enrichment in terms associated with organismal development, cardiac electrophysiology, neural function/transmission and GABA, placenta, alkaline phosphatase activity, phospholipase activity, Amyloid β, and muscle function.

DS is most commonly associated with intellectual disability [44]. Previous work has demonstrated that defects in purine metabolism and/or purine concentration in the developing brain are related to intellectual disabilities [5,8]. DS brains show decreased neuronal density by mass and volume in various brain regions (e.g. cortex, hippocampus, cerebellum), which occurs during development, possibly during gestation [45]. Purinergic signaling regulates axon guidance and growth as well as establishment of correct synaptic contacts [8] and is crucial for CNS development [46]. The GART enzyme exhibits altered spatiotemporal expression in the developing nervous system in DS. Typically, levels of GART are highly expressed in the developing cerebellum and decrease precipitously post-partum. However, in DS, GART levels persist and decrease later in development [5].

Our ClueGO results include terms related to alkaline phosphatase function, specifically differential expression of ALPP, ALPI, ALPG, and ALPL. ALPP, ALPI, and ALPG show elevated expression in crGART. ALPL, which encodes tissue non-specific alkaline phosphatase (TNAP), is elevated in HeLa. TNAP is a membrane bound extracellular enzyme present in mineralizing bones, renal tissue, and the central nervous system (CNS) [47]. In studies of murine CNS development TNAP was found to be associated with the neural tube [48] and its expression is highest in early embryonic development associating with neural precursor and progenitor cells [49]. TNAP expression has also been observed during synaptic formation and maturation [50], promoting axonal growth [51]. TNAP also has a role in the hydrolysis of extracellular nucleotides [8,52]. Extracellular ATP was found to induce migration of neural progenitor cells [53], indicating that TNAP may play a role in regulating extracellular ATP pools for migratory events. Another TNAP function is found in the metabolism of Vitamin B6, which is a cofactor for enzymes involved in neurotransmitter (e.g. GABA) synthesis [54]. Not much is known about the direct role of TNAP in proliferation and differentiation. However, given the function and spatiotemporal expression of TNAP, it is likely that TNAP is directly involved in purinergic signaling or regulation of the extracellular purine pool. Hence, altered TNAP levels may potentially be deleterious during CNS development.

Gamma-amino butyric acid (GABA) is the main inhibitory neurotransmitter in healthy adult brains and has been of particular interest in DS [45,55]. Analysis of fetal brain tissue has shown a smaller hippocampus and decreased GABA neurotransmitters in DS which suggests impaired neurogenesis or migration of GABAergic interneurons [56]. DS patients exhibit an increased incidence of epileptic seizures [57], children exhibit sleep disturbances [58] and hyperactivity [59]. These conditions may be partially due to abnormal GABA signaling. Studies employing DS murine models have shown that altered GABA signaling results in synaptic excitatory/inhibitory signal imbalance, impaired synaptic plasticity [45,55,60], and learning and memory deficits [61,62]. The enrichment for GABA-related terms within our analysis suggests that GART and DNPS may play a role in these processes.

Alzheimer’s disease (AD) is a form of dementia characterized by accumulation of neural amyloid β plaques, Tau neurofibrillary tangles, and chronic neural inflammation. Recent work supports the hypothesis that inflammation plays a critical role in AD, and therapeutic intervention designed to reduce neural inflammation shows promise. Inflammation is typically mediated by cytokine and chemokine secretion as well as fatty acid metabolism [63]. Chemokines and cytokines (such as IL-1β and TNF?) are secreted in response to injury and act as proinflammatory signals, resulting in clearance cell recruitment to the damaged tissue [63]. Fatty acid derivatives, specifically lipoxins (derived from ω-6 via phospholipaseA2 and lipoxygenase catalysis of arachidonic acid) [64] and ω-3 metabolites such as maresins, resolvins, and protectins [65] are all potent anti-inflammation mediators. ω-6 fatty acid is typically stored in the phospholipid bilayer as arachidonic acid, which is then cleaved by phospholipase activity and then metabolized via secondary enzymes to eicosanoids and lipoxins [66]. Coincidentally, extracellular ATP signaling plays a crucial role in inflammation through the purinergic P2 and P1 receptors [67]. DNPS and GART levels may play important roles in these processes.

We obtained 1282 DEGs unique to the crGART versus HeLa comparison (i.e., DEGs not found in the crADSL vs HeLa [14] and crATIC vs HeLa comparisons [15]). These DEGs mapped to Biological process ontologies related to RNA splicing and mitochondrion organization (Supplementary Table 4 - PANTHER unique). It is unclear why these genes are significantly changed only in the crGART cell line. One possibility is that the absence of GART may have an effect unrelated, or peripherally related, to DNPS that affects the transcription of these genes.

In conclusion, our results indicate that DNPS deficiency affects the cellular transcriptome, significantly altering the expression of over 4000 genes. We believe that the crGART cell line is an attractive model of DNPS deficiency in the absence of substrate accumulation and will be a valuable tool for investigating the role of DNPS dysregulation in DS and other disorders.

Further, given the remarkable difference in CD36 transcript levels and protein expression in crGART versus Hela, as well as the difference in LCFA levels and rescue of CD36 expression in crGART versus transfected crGART (crGART/K1-GART), we believe that the cell line will be a good working model for investigating CD36 function in the context of DNPS and purine metabolism.

## Supporting information

Supplementary Table 1 - DEGs

Supplementary Table 2 - ClueGO

Supplementary Table 3 - PANTHER

Supplementary Table 4 - PANTHER unique

Supplementary Table 5 - PANTHER common

Supplementary Table 6 - Metabolon LCFA

## Abbreviations

(APRT): adenine phosphoribosyltransferase
(AMP): adenosine monophosphate
(ATP): adenosine triphosphate
(ADSL): adenylosuccinate lyase
(AD): Alzheimer disease
(ZMP): 5-aminoimidazole-4-carboxamide ribonucleotide
(ATIC): 5-aminoimidazole-4-carboxamide ribonucleotide formyltransferase / inosine monophosphate cyclohydrolase
(AMPD): AMP deaminase
(CNS): central nervous system
(DNPS): de novo purine synthesis
(DEG): differentially expressed gene
(DS): Down syndrome
(FDR): false discovery rate
(FCM): fetal calf macroserum
(FCS): fetal calf serum
(GABA): gamma-amino butyric acid
(GO): gene ontology
(GAR): glycineamide ribonucleotide
(GMP): guanosine monophosphate
(IMP): inosine monophosphate
(5-PRA): phosphoribosylamine
(PAICS): phosphoribosylaminoimidazole carboxylase / phosphoribosylaminoimidazole succinocarboxamide synthetase
(GART): phosphoribosylglycinamide formyltransferase / phosphoribosylglycinamide synthetase / phosphoribosylaminoimidazole synthetase
(PFAS): phosphoribosylformylglycinamidine synthase
(PRPP): phosphoribosyl pyrophosphate
(PRAT): pyrophosphate amidotransferase
(XMP): xanthine monophosphate

## Acknowledgements

The Cancer Center and the Genomics (Microarray) Shared Resource are supported in part by the Cancer Center Support Grant #P30-CA046934 from the National Cancer Institute.

MZ and VB were supported by Charles University [programmes PRIMUS/17/MED/6 and PROGRES Q26/LF1] and by the Ministry of Education, Youth and Sports of CR [LQ1604 National Sustainability Programme II].

## Author Contributions

R.M., D.P., and G.V. designed the experiments. R.M. performed the ClueGO analyses and was primary author of the manuscript. V.B. and M.Z. created and provided the CRISPR-Cas9 GART KO HeLa cell line, crGART. N.D. performed HPLC-EC analysis to detect DNPS metabolites and intermediates. T.G.W. provided sample preparation expertise. G.V. performed salmon and DESeq2 processing of fastq sequence and provided informatics support. G.V. and D.P. directed and supervised the team members and manuscript preparation. All authors reviewed and edited the manuscript.

