## Supplementary Figures 1-5 for "Transcriptome and metabolome analysis of crGART, a novel cell model of de novo purine synthesis deficiency: Alterations in CD36 expression and activity"

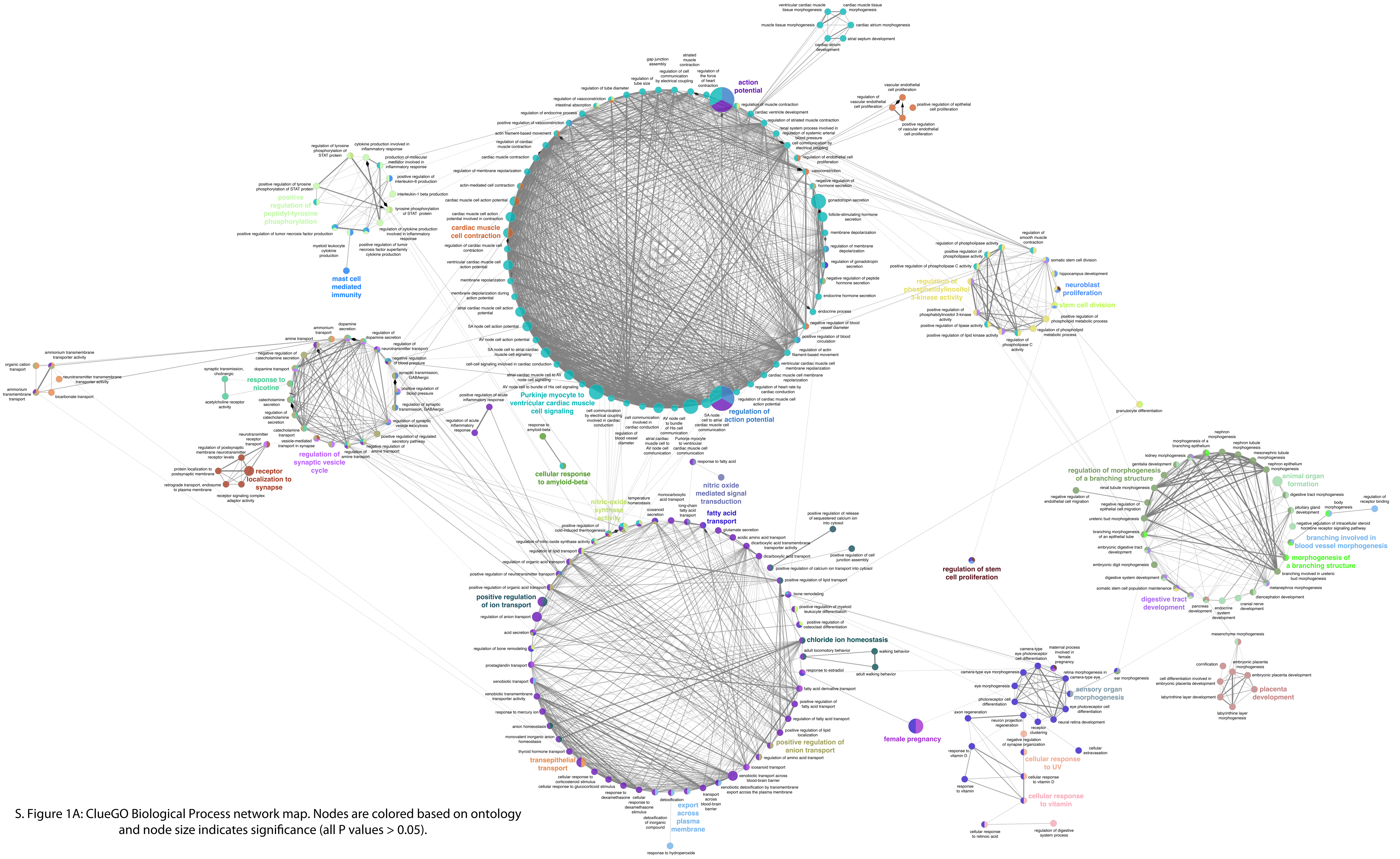


S. Figure 1A: ClueGO Biological Process network map. Nodes are colored based on ontology and node size indicates significance (all P values > 0.05).

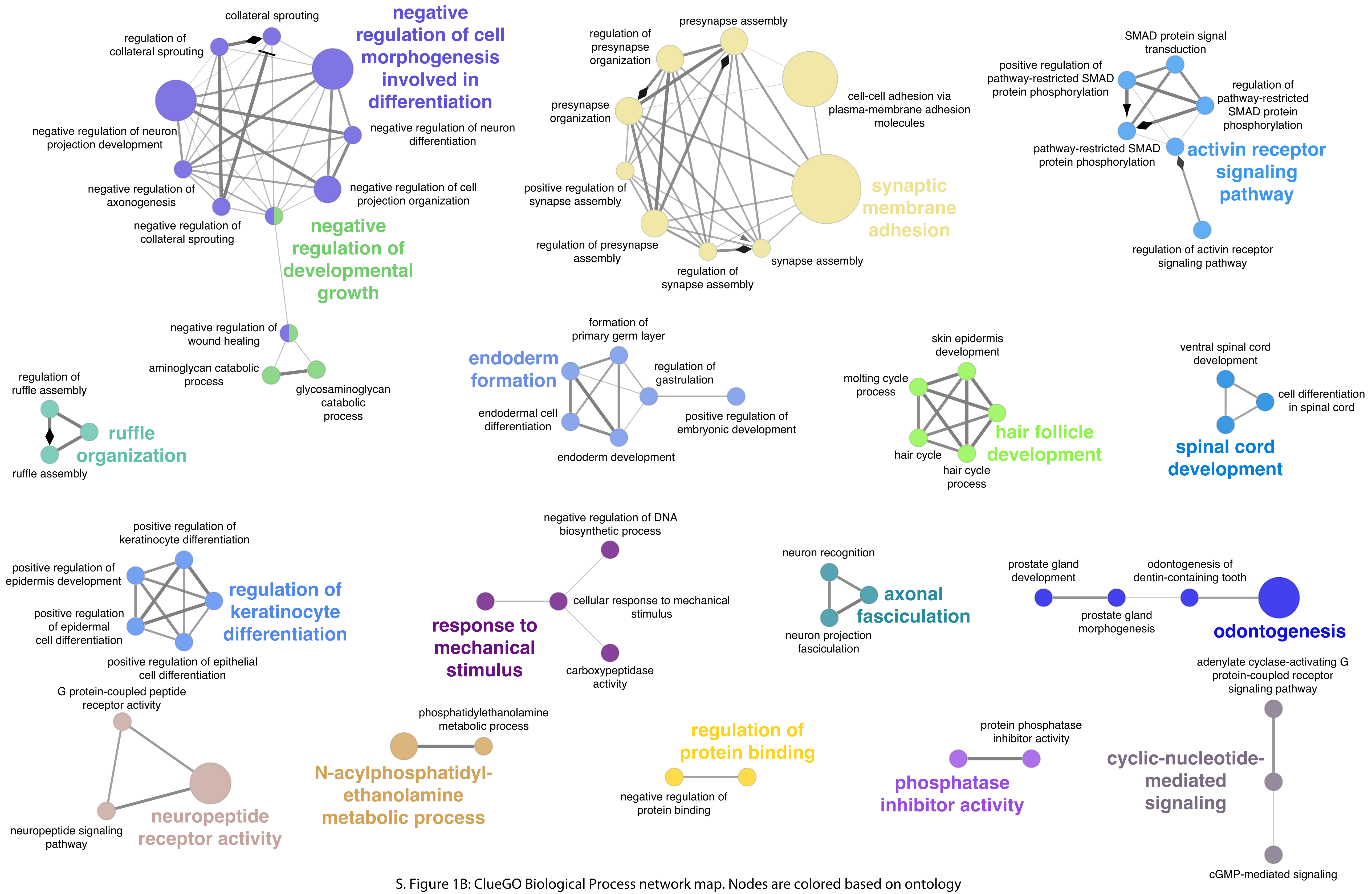


S. Figure 1B: ClueGO Biological Process network map. Nodes are colored based on ontology and node size indicates significance (all P values > 0.05).

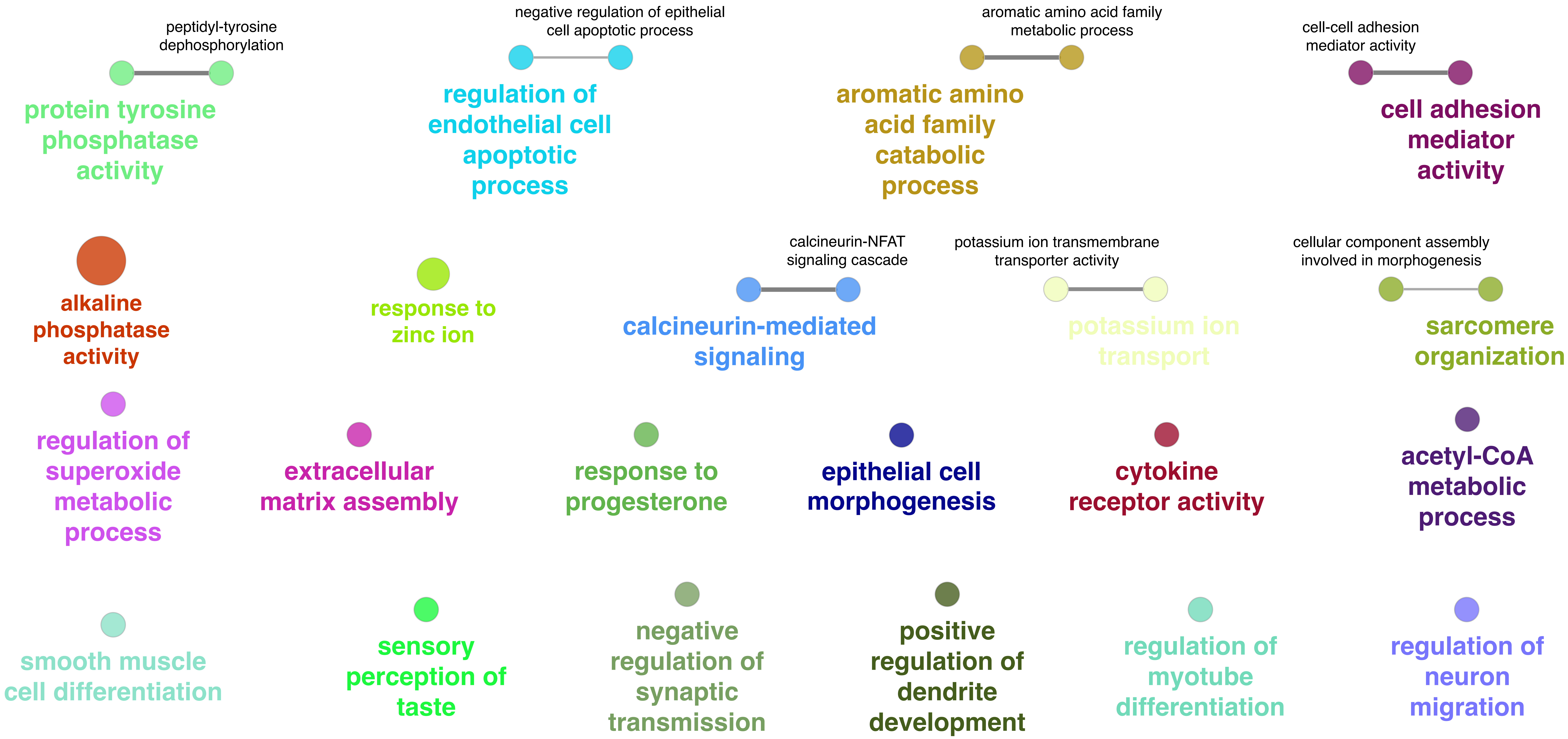


S. Figure 1C: ClueGO Biological Process network map. Nodes are colored based on ontology and node size indicates significance (all P values > 0.05).

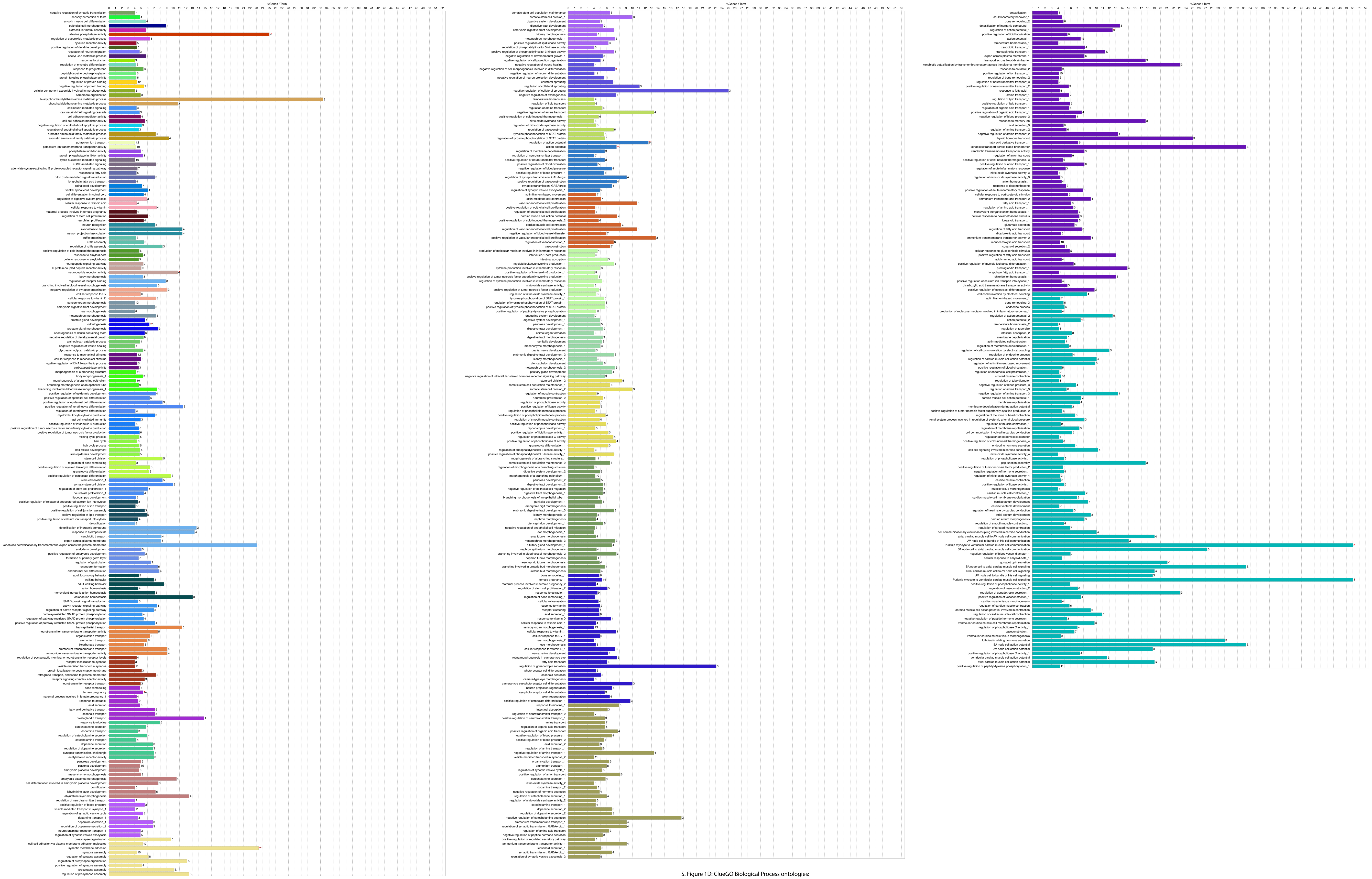


S. Figure 1D: ClueGO Biological Process ontologies:
Number and percent genes per ontology

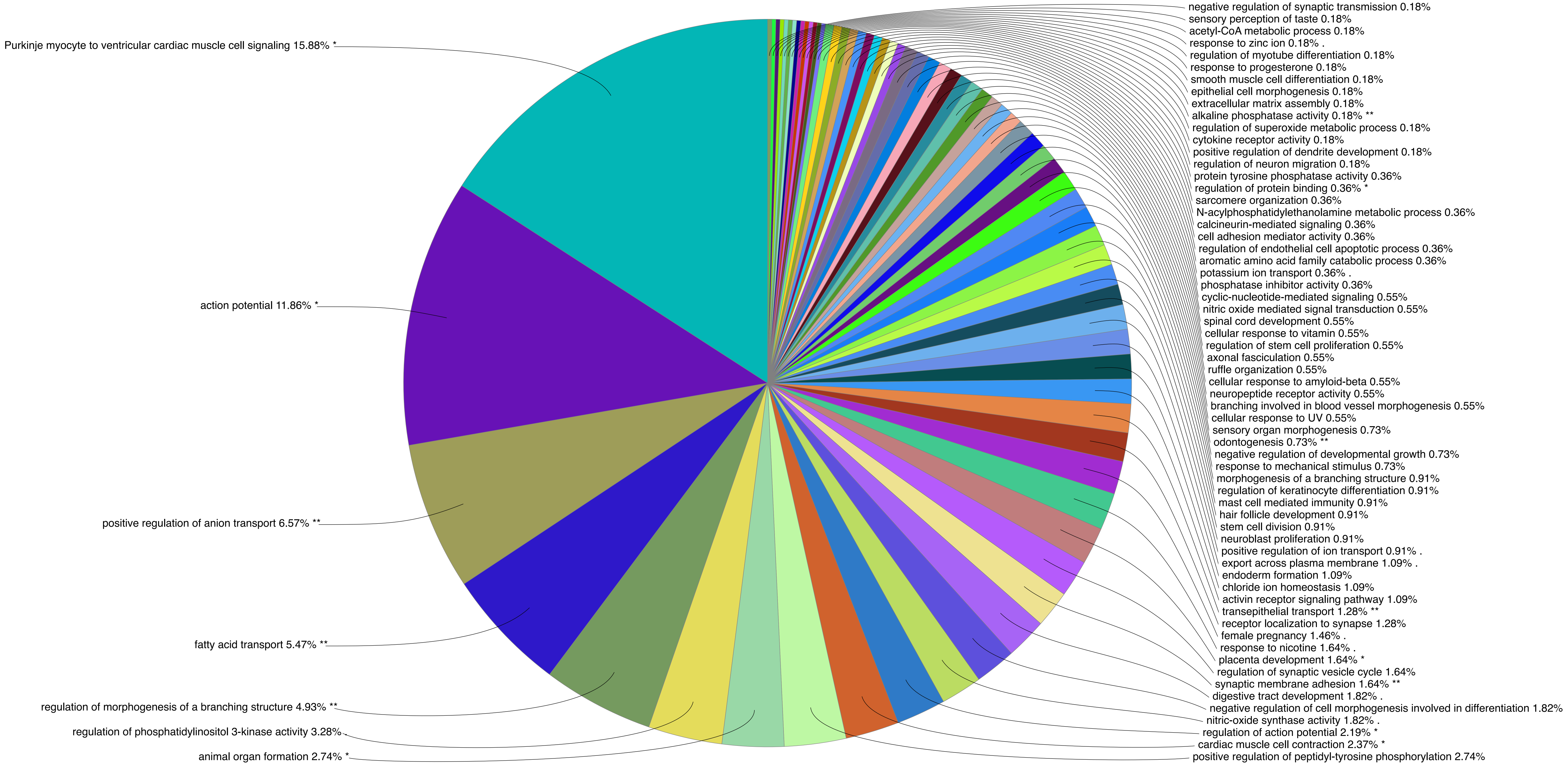


S. Figure 1E: ClueGO Biological Process ontologies, percent representation:
Colors represent ontology groups.

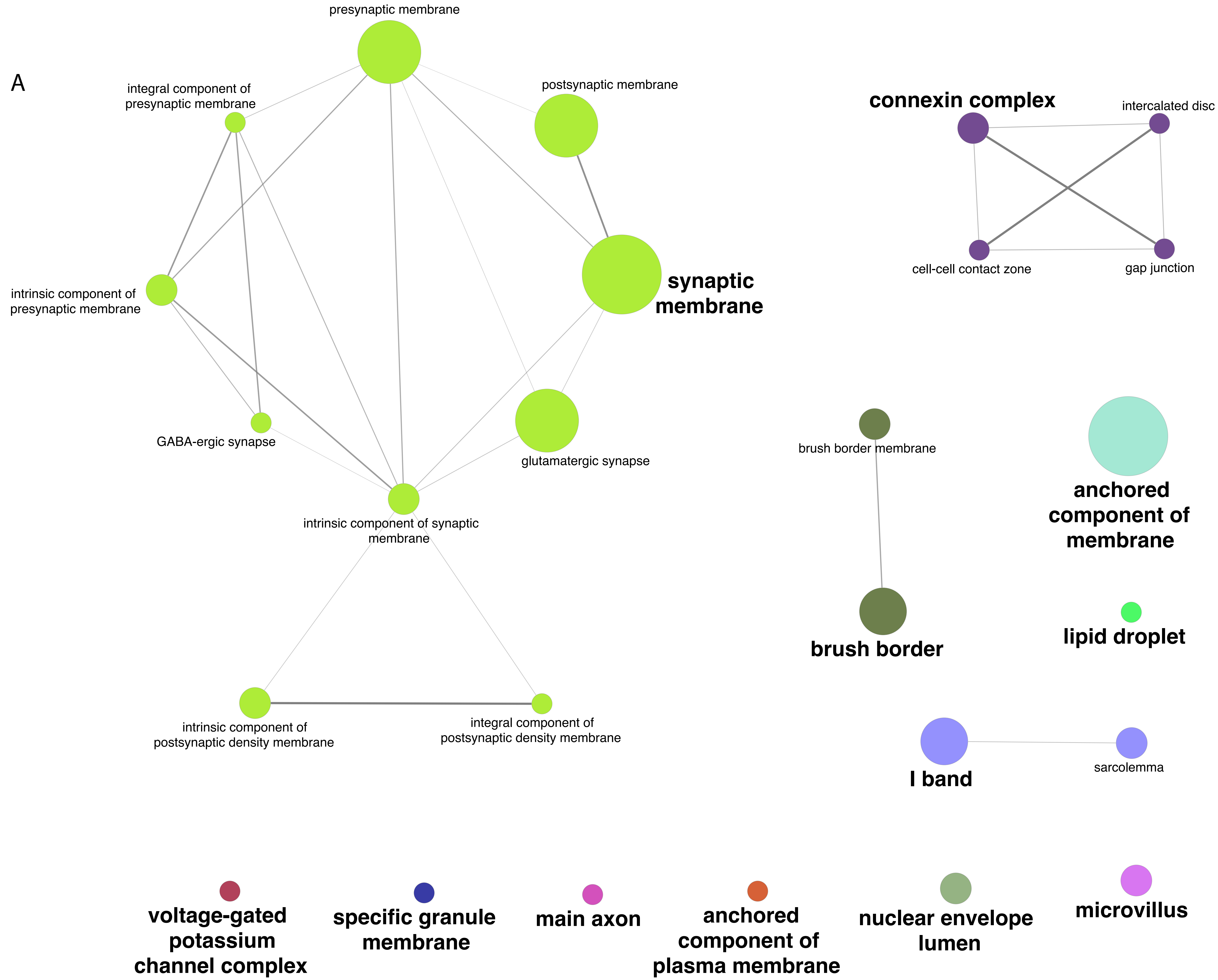


S. Figure 2A: ClueGO Cellular Component network map. Nodes are colored based on ontology and node size indicates significance (all P values > 0.05).

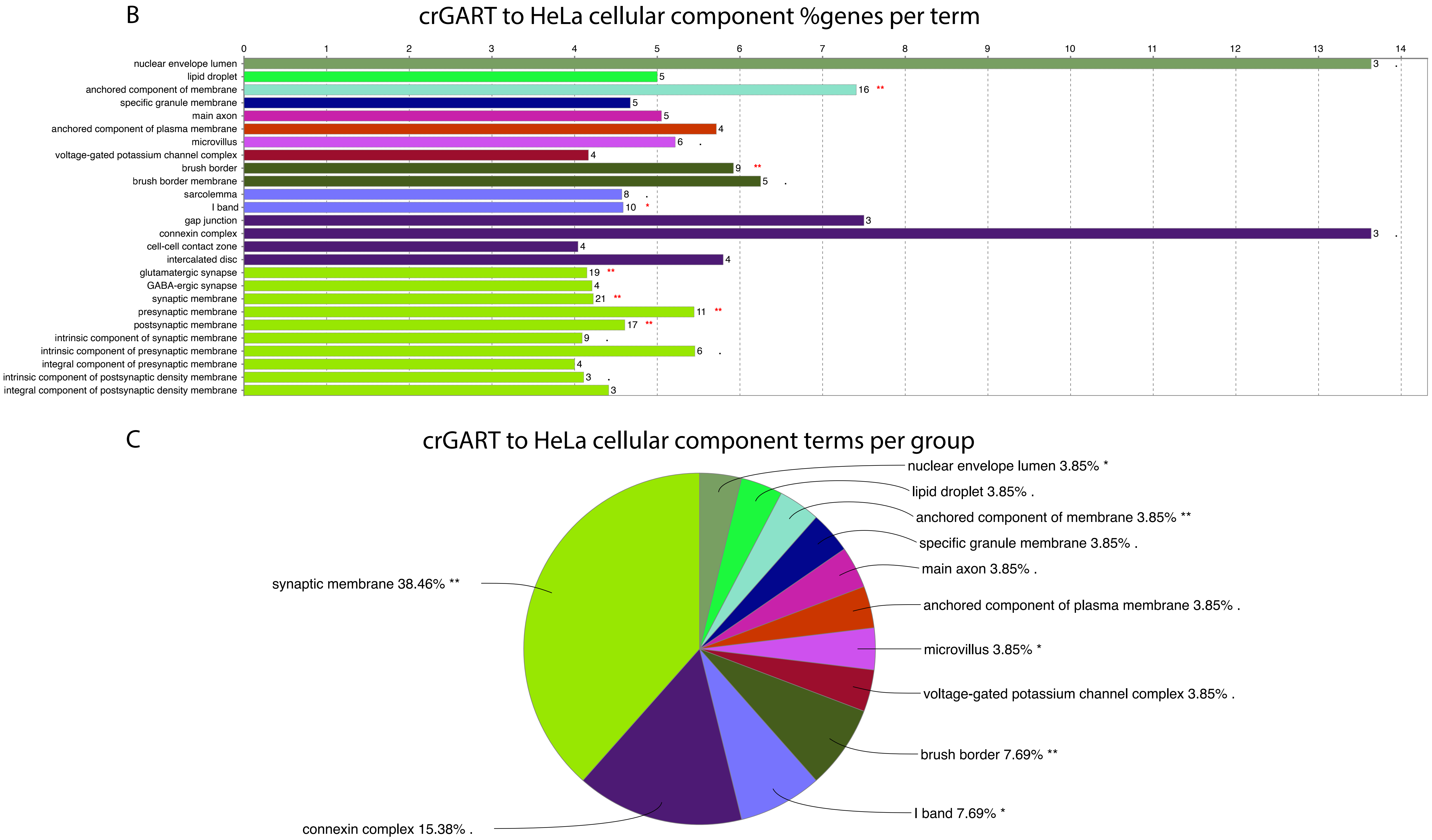


S. Figure 2: B) ClueGO Cellular Component ontologies: Number and percent genes per ontology
C) Percent representation: Colors represent ontology groups.

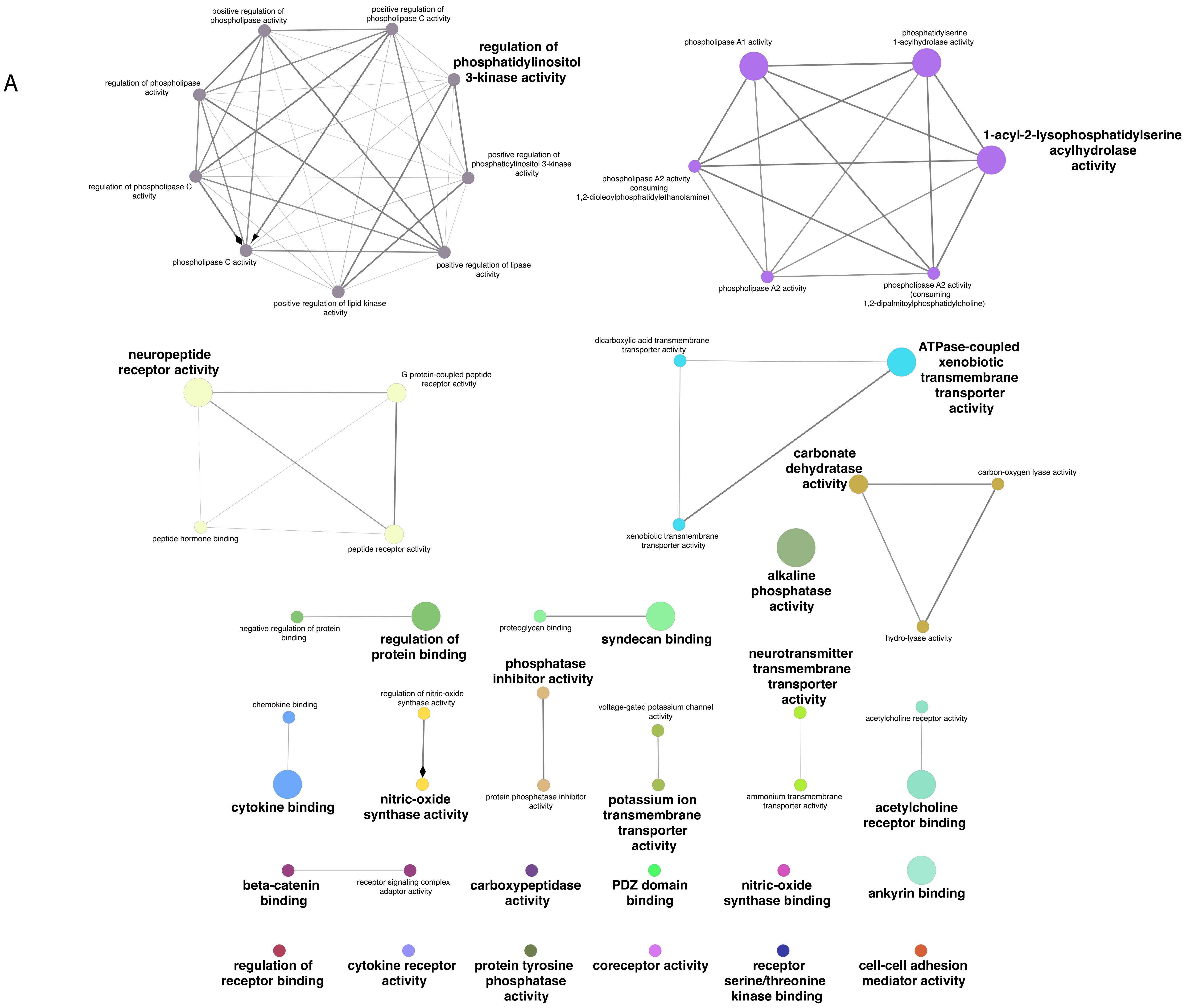


S. Figure 3A: ClueGO Molecular Function network map. Nodes are colored based on ontology and node size indicates significance (all P values > 0.05).

B

### crGART to HeLa molecular function %genes per term

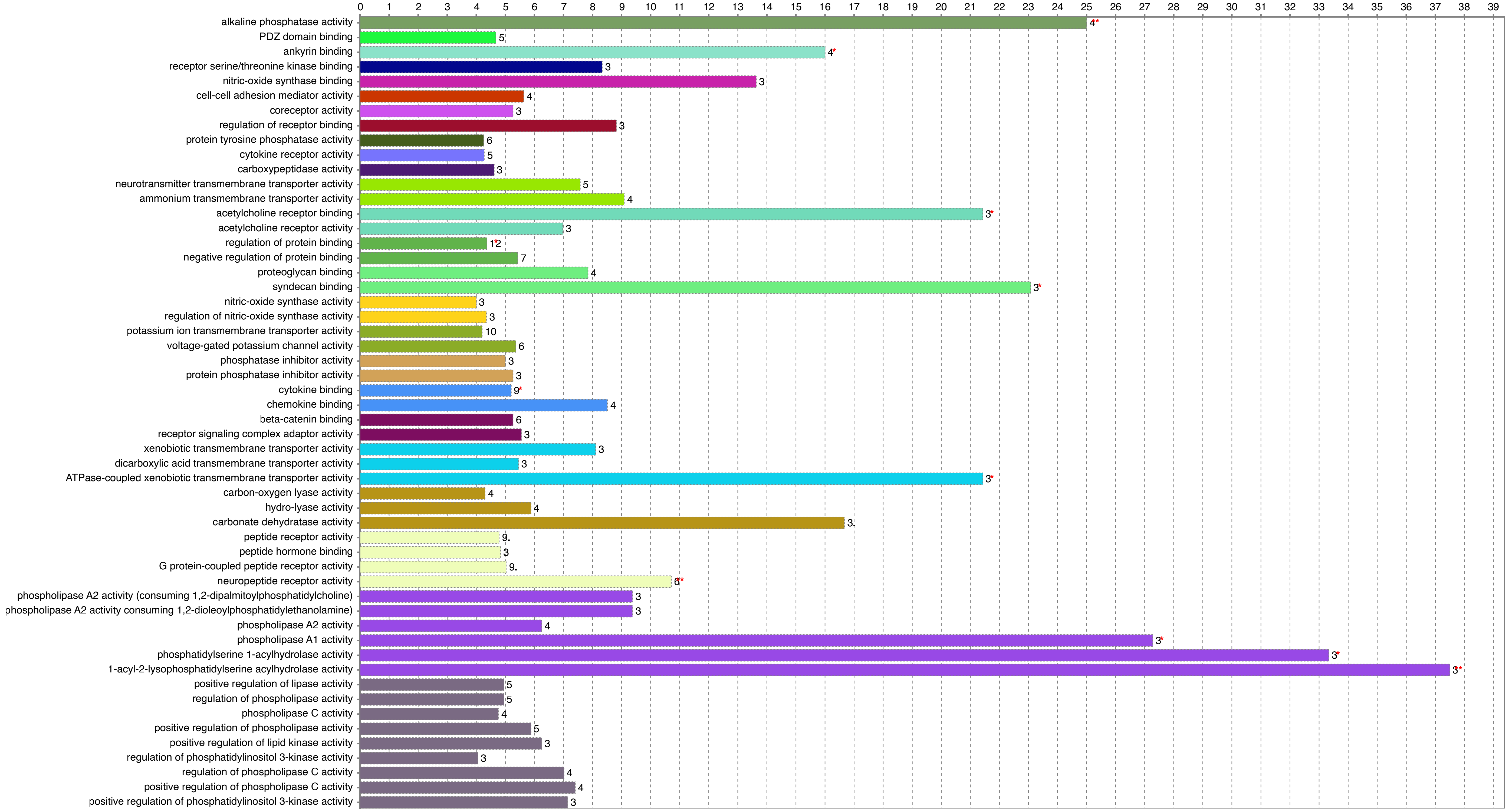


C

### crGART to HeLa molecular function terms per group

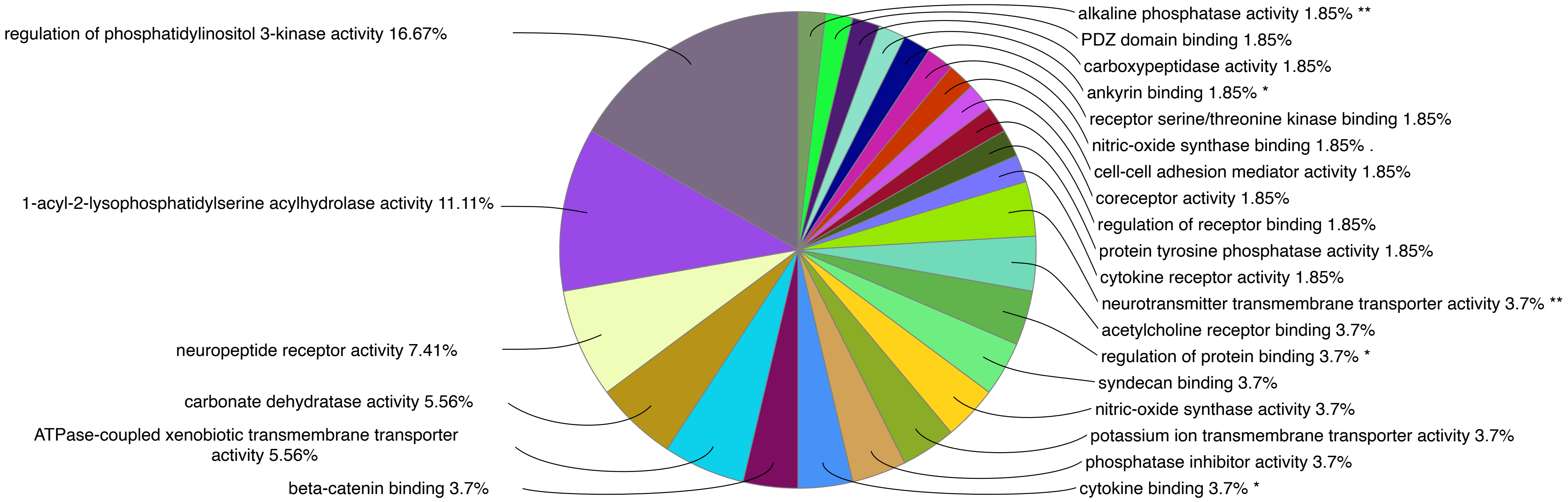


S. Figure 3: B) Molecular Function ontologies: Number and percent genes per ontology
C) Percent representation: Colors represent ontology groups.

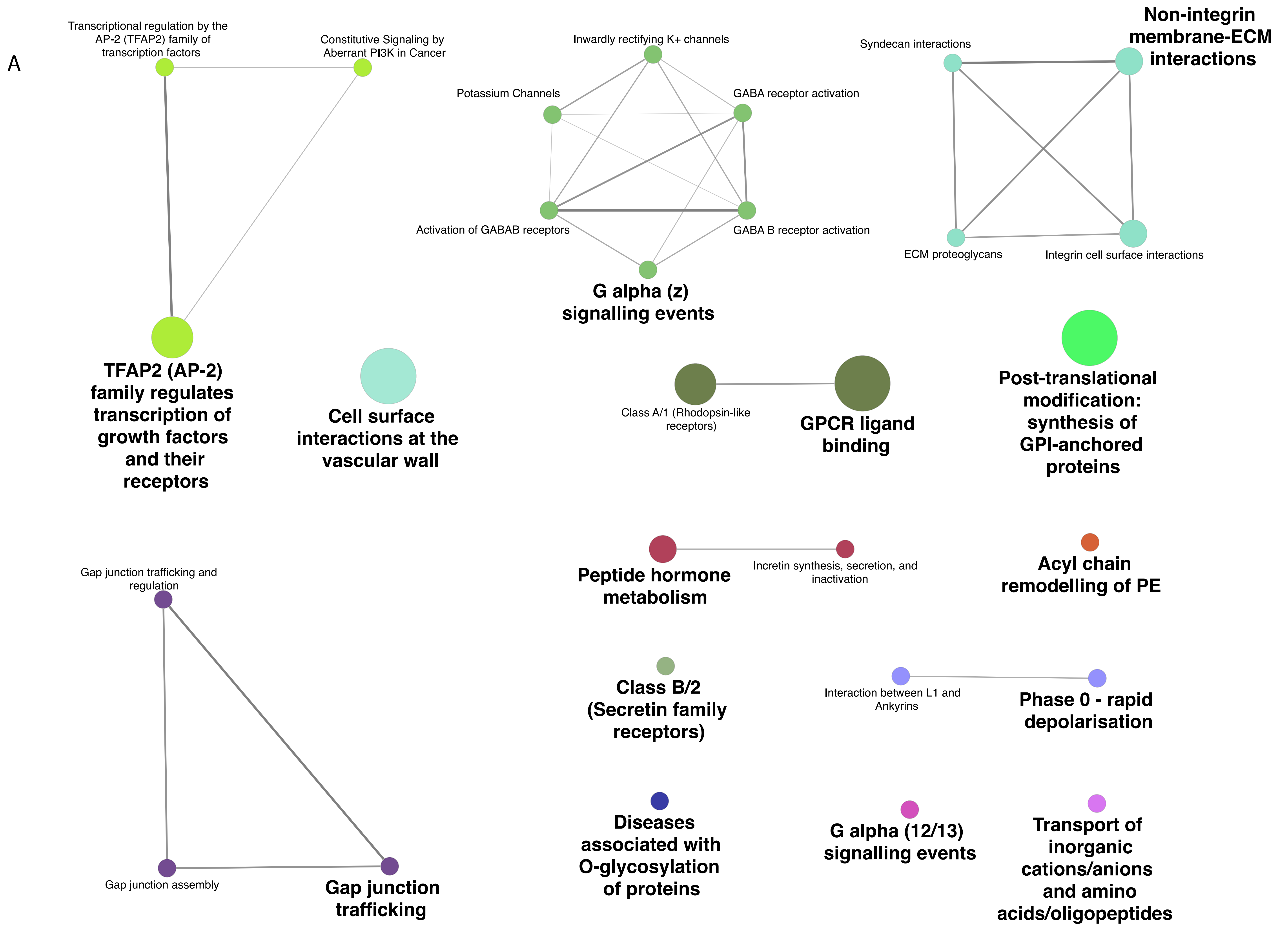


S. Figure 4A: ClueGO Reactome Pathways network map. Nodes are colored based on pathway and node size indicates significance (all P values > 0.05).

B

### crGART to HeLa Reactome pathways %genes per term

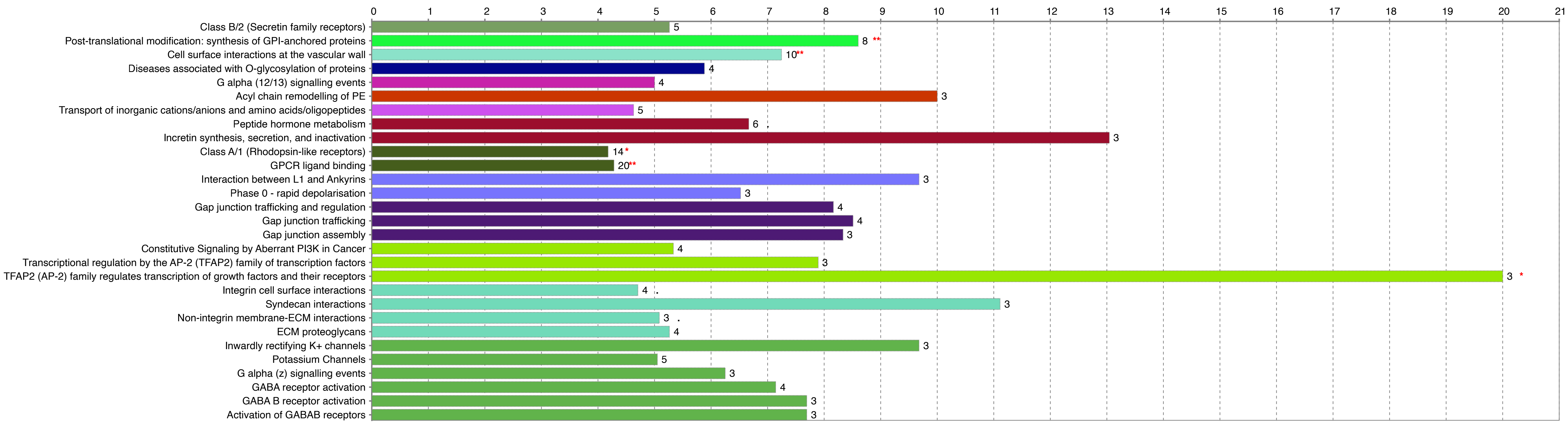


C

### crGART to HeLa Reactome pathways terms per group

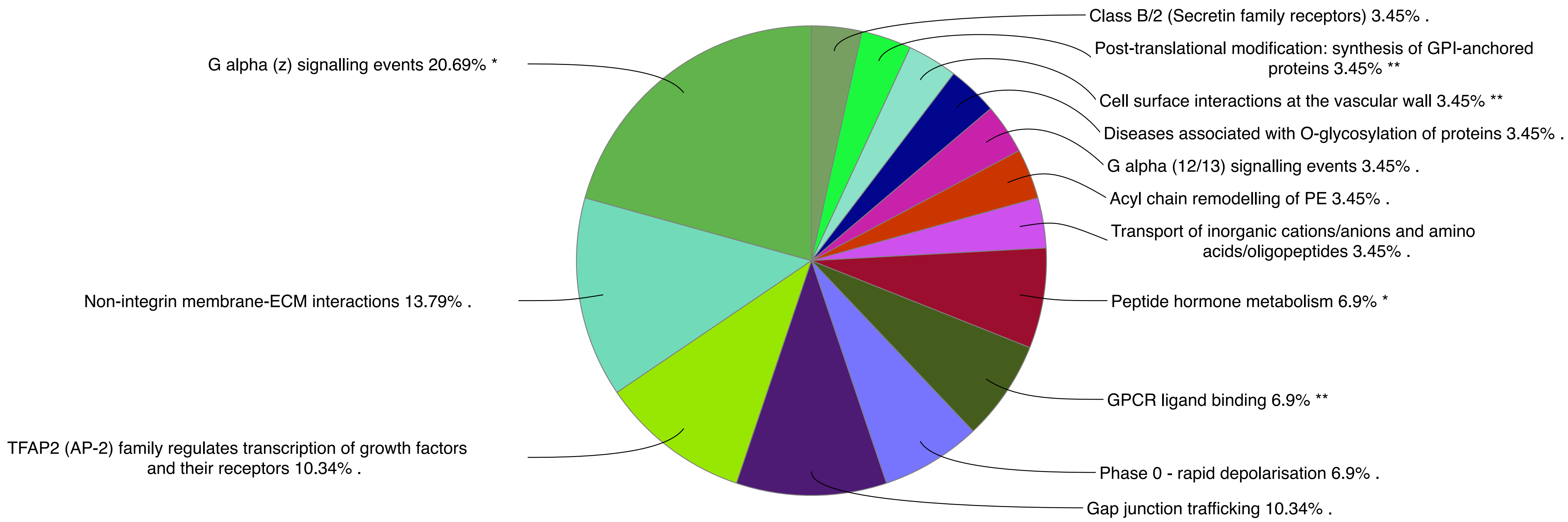


S. Figure 4: B) Reactome Pathways: Number and percent genes per pathway C) Percent representation: Colors represent pathway groups.

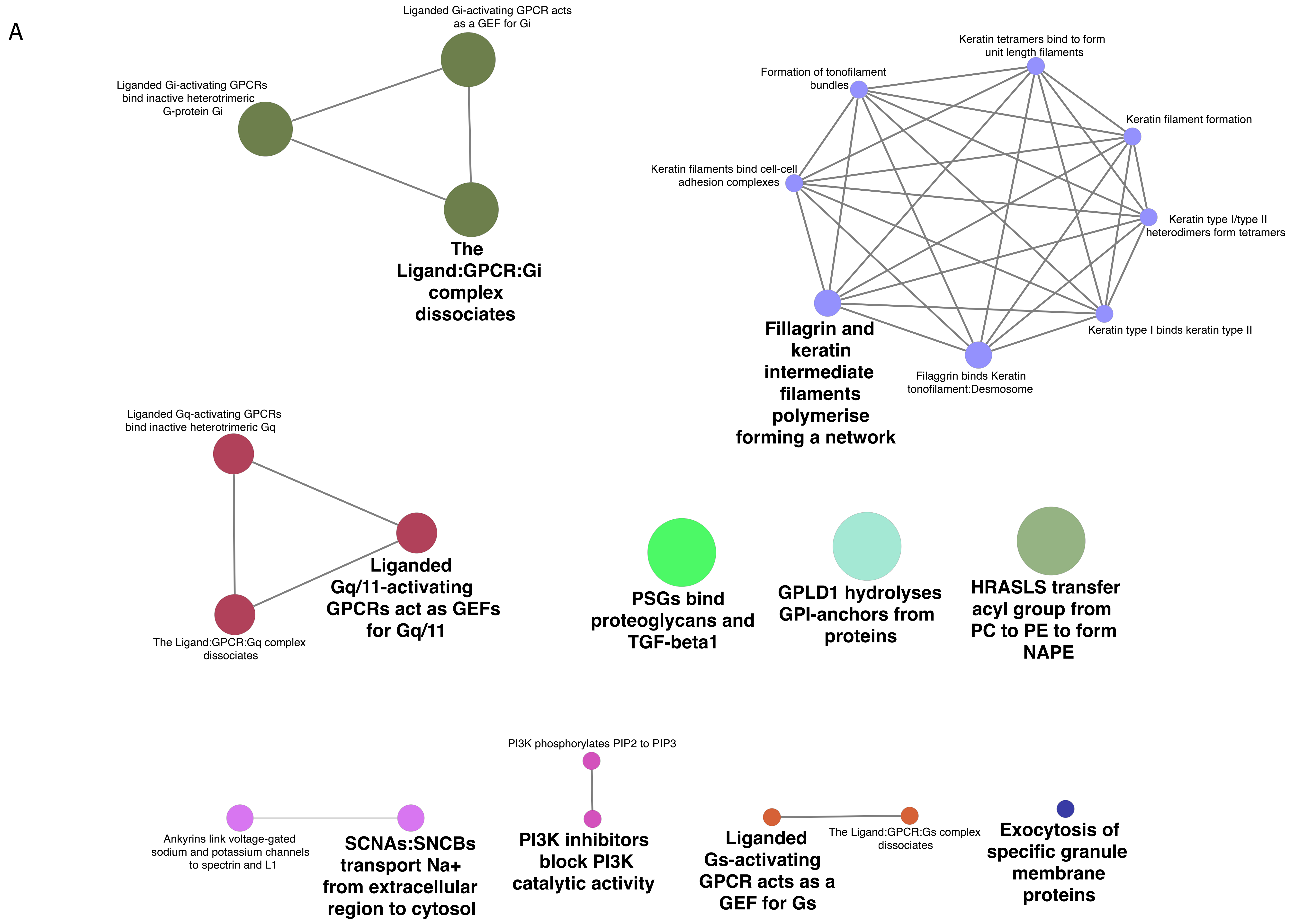


S. Figure 5A: ClueGO Reactome Reactions network map. Nodes are colored based on reaction and node size indicates significance (all P values > 0.05).

B

### crGART to HeLa Reactome reactions %genes per term

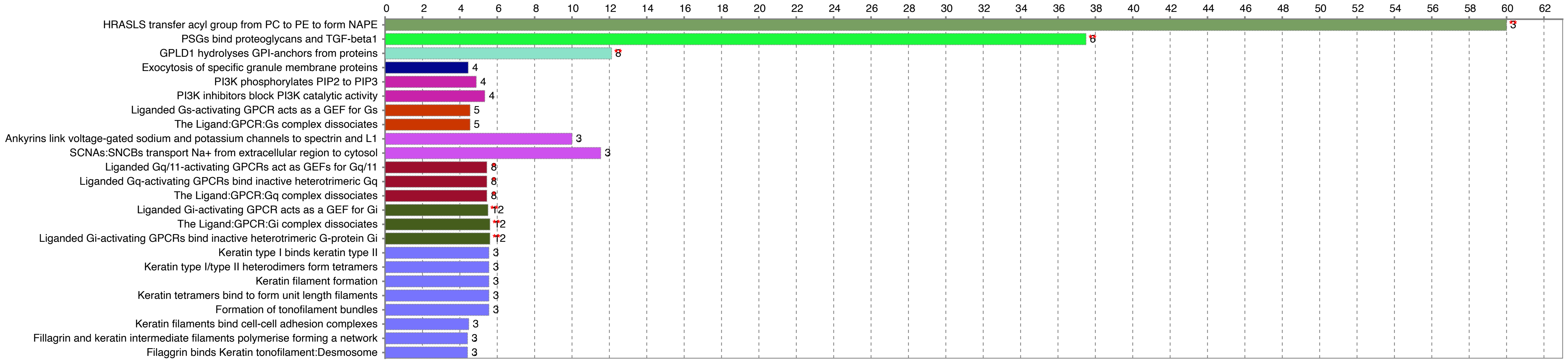


C

### crGART to HeLa Reactome reactions terms per group

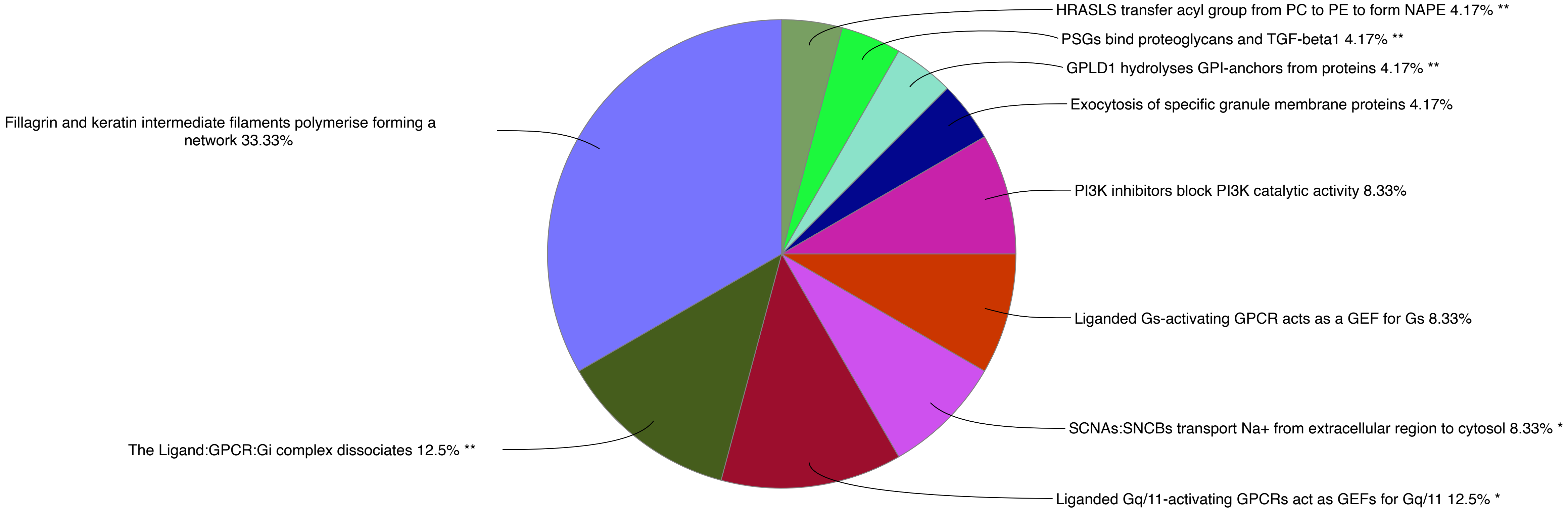


S. Figure 5: B) Reactome Reactions: Number and percent genes per reaction C) Percent representation: Colors represent reaction groups.
